## Supplementary Figures and Tables for "Subcellular mass spectrometry imaging of lipids and nucleotides using transmission geometry ambient laser desorption and plasma ionisation"

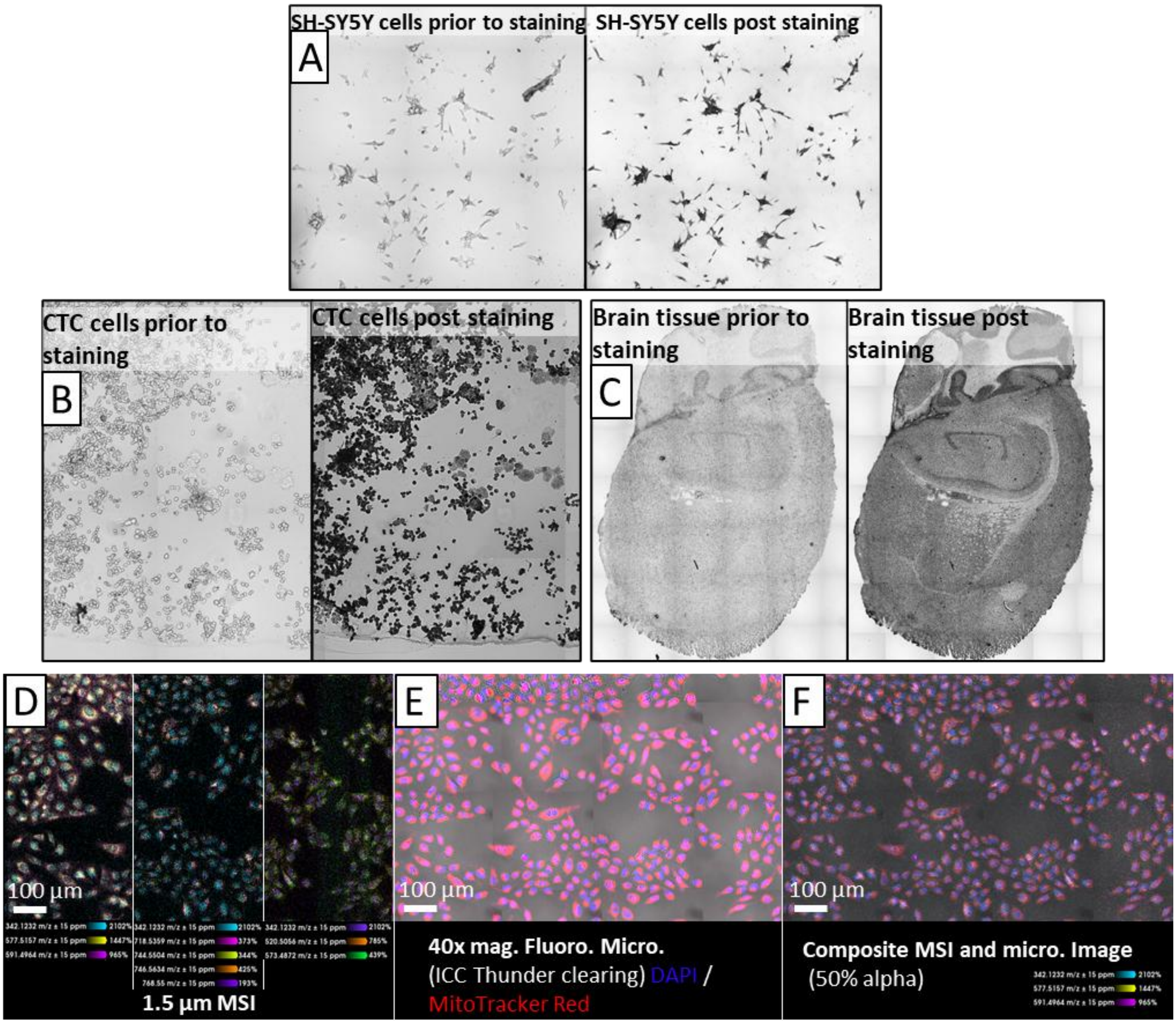

**Supplementary Fig. 1: Effect of cresyl violet (CV) pre-staining on analyte delocalisation and sample displacement.**

**(A)** Left-to-right: 10x brightfield microscopy of SH-SY5Y cells prior to staining, 10x brightfield microscopy of SH-SY5Y cells post CV staining, and 10x brightfield microscopy of SH-SY5Y cells post CV staining. The data series displays that the spacing between cells before and after staining is spatially maintained, and no loss of adhered cells has occurred. **(B)** Replicate experiment of panel (A), showing that the sample spacing, and cell adherence is also observed to be unaffected for a plated circulating tumour cell (CTC) line. Larger particulates, presumed to be artifacts leftover from cell media, are also unaffected and can be observed in before and after staining micrographs. **(C)** Displays 10x brightfield microscopy from a mouse brain section taken before and after CV staining. Any defects to the tissue section (*e.g.,* holes in the tissue, fraying around the edges) is observed to be present in the tissue prior to staining being undertaken. **(D-F)** CV pre-stained U2OS cells imaged using 1.5 µm MSI and displaying a total of 9 overlaid ion intensity images split across separate thirds of the MSI (D), 40x fluorescence and brightfield microscopy (E), and a composite of both images using 50% transparency. As can be observed for the 9 analytes being displayed in the MSI (E) and the co-registered composite image (F), analyte signal is confined to 'on cell' regions with minimal-to-no analyte signal being observed off the cells.

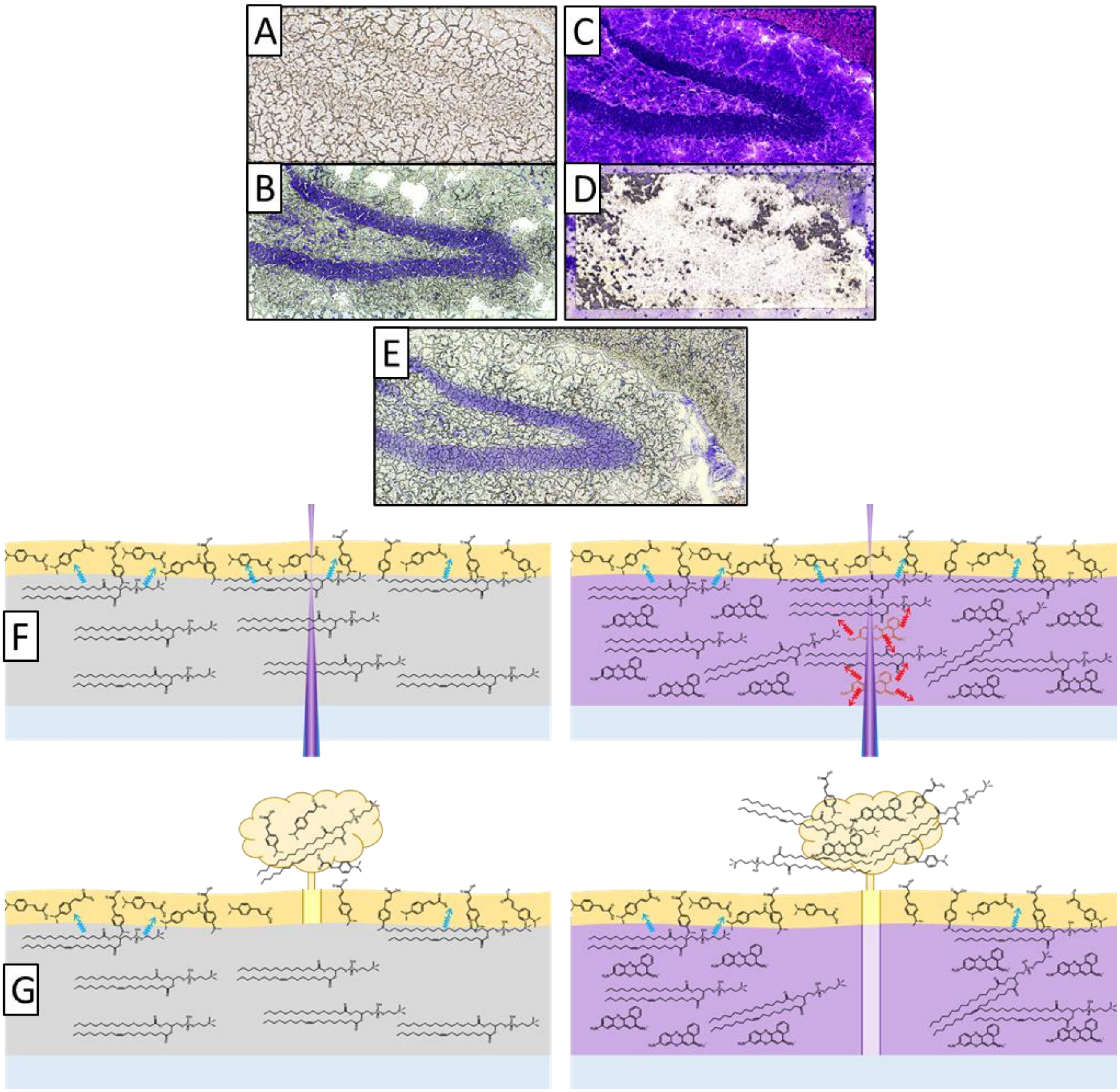

**Supplementary Fig. 2: Examination of tissues pre and post MSI ablation, with or without CV pre-staining.**

**(A)** 10x brightfield microscopy of a murine hippocampus without CV pre-staining and before MSI. **(B)** 10x brightfield microscopy of a murine hippocampus with CV pre-staining and before MSI. **(C)** 10x brightfield microscopy of a murine hippocampus without CV pre-staining and after MSI. Prior to this micrograph, the sample was de-lipidated and stained with CV using conventional methods indicated in the Methods section. The edge of the ablation region can vaguely be discerned as a light discolouration to the tissue. Additionally, holes within the ablation region can be identified, but cannot be pinpointed to deformations occurring during the MSI or histological staining methods. **(D)** 10x brightfield microscopy of a murine hippocampus with CV pre-staining and after MSI. The edge of the ablation region can be easily discerned, and objectively large volumes of tissue are absent from the region that had been illuminated by the laser. **(E)** 10x brightfield microscopy of a murine hippocampus that serves as a reference point for comparison against the other micrographs. This tissue has undergone was not exposed to laser ablation and has only undergone conventional CV staining methods prior to microscopy. **(F-G)** A diagram displaying a representation of how the presence of CV is affecting laser ablation events. Panel (F-left) displays a cross sectional view of a tissue (grey) containing lipids, mounted on a glass microscopy slide (light-blue), coated in a photoactive matrix (yellow). The lipids from the tissue form co-crystals with the matrix layer (blue arrows). Panel (F-right) displays the same cross-sectional view, but instead the tissue is stained purple from the presence of CV throughout. Within both panels of (F), the transmission laser is focussed through the rear of the sample slide and into the surface matrix layer. When the tissue does not contain CV, the laser light transmits through the non-photoactive tissue into the matrix layer, leading to ejection of matrix-lipid co-crystals and the formation of a MALDI plume (G-left). Instead, when CV is present within the tissue, the CV molecules in the path of the laser beam are excited and release heat to the surrounding tissue material (F-right). This leads to the ejection of a ‘column’ of tissue material in the laser beam path as well as the matrix layer (G-right).

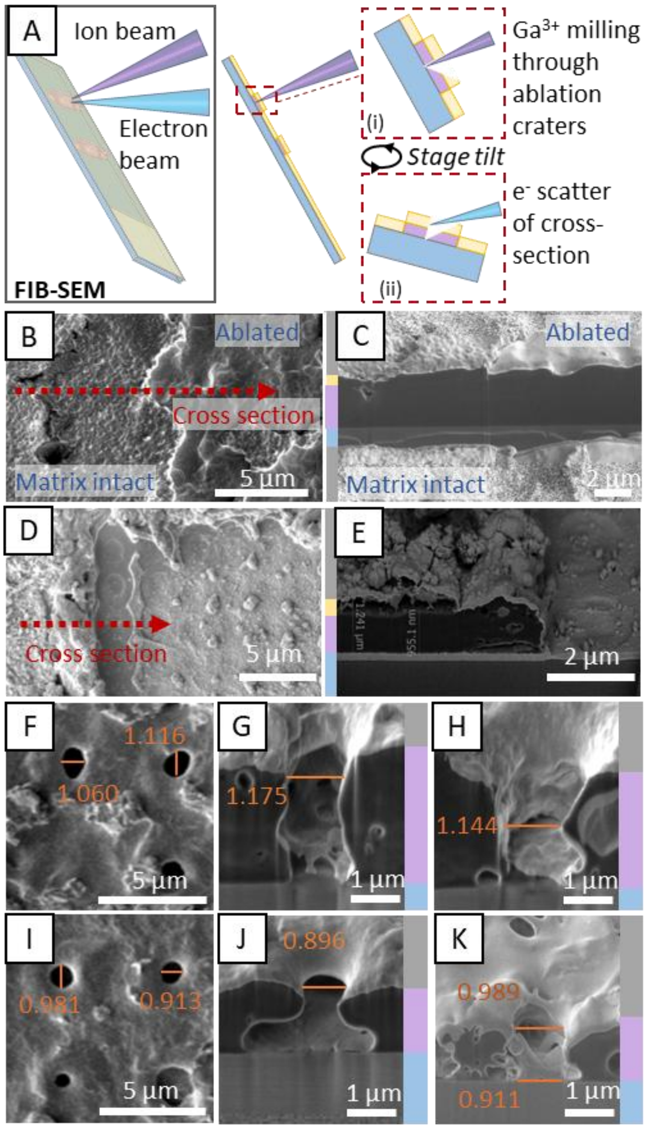

**Supplementary Fig. 3: Focused ion beam-scanning electron microscopy (FIB-SEM) validation of t-MALDI ablation crater size and form.**

**(A)** Schematic displaying FIB-SEM mechanics. After surface SEM is obtained, a focussed beam of $Ga^{3+}$ is used to mill through the matrix + sample layers. The stage is then tilted by 52°, allowing for SEM visualisation of the newly created cross-section. **(B-C)** 2.5 µm ablation array of sublimed DMACA matrix with no cresyl violet staining of the tissue layer. (B) 2500× SEM shows the rough surface of matrix layer still intact and the smoother surface after it has been ablated away by NUV laser. (C) 5000× FIB-SEM shows a cross section at the ablation array boundary, revealing the tissue layer appears unperturbed by NUV laser transmitting through, while the surface matrix is absent. **(D-E)** 2.5 µm ablation array of sublimed DMACA matrix with additional cresyl violet staining of the tissue layer. (D) 2500× SEM shows the matrix and tissue layers are still intact, while significant material loss is observed after it has been ablated by NUV laser. (E) 25000× SEM shows a cross section at the ablation array boundary, revealing the increased sampling volume with the presence of cresyl violet. **(F-H)** 5 µm ablation arrays (~1 µm craters) from DHA sublimed tissues with cresyl violet staining. (F) Displays 800× SEM images of the arrays surface. (G-H) Displays the 5000× SEM images through the FIB generated cross-sections of the ablation array. **(I-J)** 5 µm ablation arrays (~1 µm craters) from DMACA sublimed tissues with cresyl violet staining. (I) Displays 800× SEM images of the arrays surface. (J-K) Displays the 5000× SEM images of the FIB generated cross-sections through the ablation array. To assist interpretation, colour bars to the right of these images show the layers corresponding to the glass slide (blue), the tissue layer (purple), the matrix layer (yellow; if discernible) and the surface layer tilted into the page (grey). All annotation values are in µm.

**(Extended text for Supplementary Fig. 3) Ablation spot size and crater dynamics.**

A nuance of a transmission geometry MALDI source is the laser initially transmitting through the sample before entering the matrix layer. As the plasma post ionisation device allows for decoupling of the MALDI desorption and ionisation events, we are able to exploit this optical geometry by including an additional photo absorbing matrix in the tissue layer (*cf.* main text Fig. 2). This allows us to increase the ablation volume per pixel area and significantly improve ion yields for MSI. To investigate ablation crater dynamics and laser focal spot size, scanning electron microscopy (SEM) employing focused ion beam (FIB) milling was obtained for MALDI ablation arrays on murine brain tissue (*cf.* Supplementary Fig. 3).

Focussed ion beam-scanning electron microscopy (FIB-SEM) uses a fine beam of ions, such as $Ga^{3+}$ $3^{+}$ used here, to mill through a sample and create cross sections. By tilting the sample stage to access this milled surface, the electron beam can be focussed on the cross section and secondary electrons and electron back-scatter can detected through conventional electron microscopy methods (*cf.* Supplementary Fig. 3A schematic). We applied this technique to 2.5 µm ablation craters laterally spaced by 2.5 µm, which were generated using the t-MALDI-P source on DMACA sublimed mouse brain sections either having undergone cresyl violet pre-staining or not. When the tissue was left unstained, SEM imaging of the ablation array perimeter revealed a smooth surface in the areas irradiated by the laser (*cf.* Supplementary Fig. 3B; right). FIB milling through the centre of a row in the array confirmed that the matrix had been ablated from the surface, however the underlying tissue layer was left largely intact suggesting minimal interaction between the laser and tissue material (*cf.* Supplementary Fig. 3C). In contrast, the CV pre-staining of the tissue led to significant ablation of the tissue material, which is visible through both the surface SEM (Supplementary Fig. 3D) and FIB-generated cross section (Supplementary Fig. 3E) images. To assist interpretation of these cross sections, coloured bars have been included towards the edge of these images, which depict the cross-sectional layers corresponding to the glass slide (blue), the tissue layer (purple), the matrix layer (yellow; if discernible) and the surface layer (grey), which extends into the page due to the SEM stage tilting (refer to Supplementary Fig. 3A schematic).

By lowering the laser pulse energy applied in the t-MALDI experiment, smaller ablation craters of ~1 µm could be generated for pre-stained brain tissues coated with either DHA (Supplementary Fig. 3F-H) or DMACA (Supplementary Fig. 3I-K). While validating that the 1 µm MS images in main text Figure 3 are not – or are only slightly – in oversampling conditions, these lowered laser energies also allowed us to observe how the inclusion of cresyl violet into the tissue may be improving sampling volume. Of note, the ablation crater cross section images (Supplementary Fig. 3G-H, 3J-K) reveal small cavitations that surround the ablation craters, which we hypothesise are being formed by photo excitation of the cresyl violet molecules and subsequent thermal transfer and expansion. This process is advantageous as it allows for ejection of analytes from within the tissue and not solely from desorption of the analyte-matrix co-crystals. To the best of our knowledge this represents the first time the FIB-SEM technique has been used to explore MALDI ablation crater dynamics and shows promise for further probing of photo absorbers for improved detection sensitivity in high spatial resolution MSI experiments and the fundamental physics processes occurring in t-MALDI.

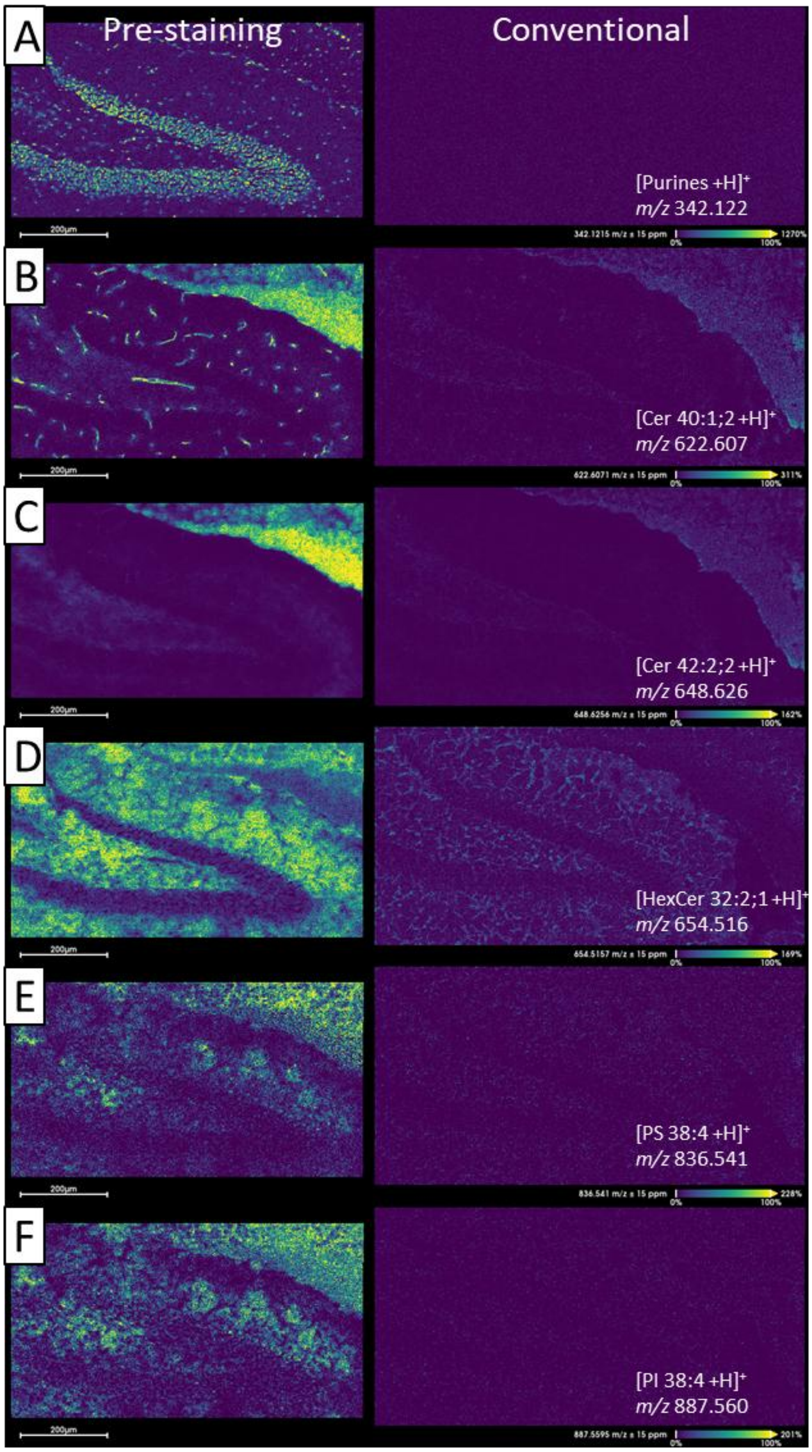

**Supplementary Fig. 4: Comparison of individual ion intensity distributions used in Main text Fig. 2 MSI overlays.**

Panels A-F are divided into two columns, with the left displaying 2 µm MSI of the pre-stained hippocampus and the right displaying the 2 µm MSI from the conventionally prepared hippocampus. Each image pair is of the lipid *m/z* indicated and share the same ion intensity scale. Across the wide array of lipid classes displayed, pre-staining is observed to outperform conventional preparations in terms of the ion intensity and the overall spatial information available.

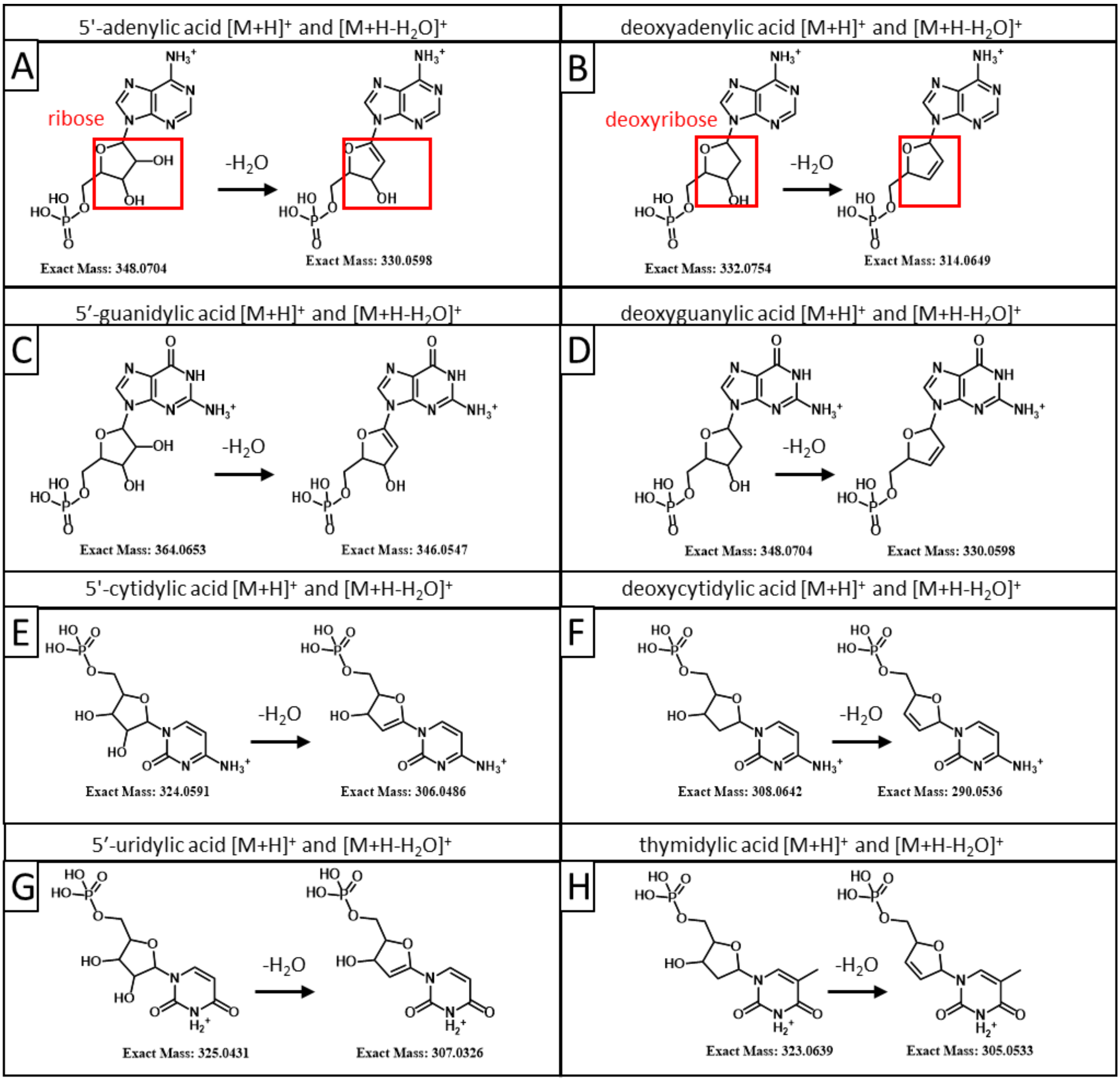

**Supplementary Fig. 5: Molecular ion structures of nucleotides observed in AP-MALDI-P.**

While all indicated *m/z* values can be observed within tissue and cell experiments, the dehydrated cation $[M+H-H_2O]^+$ is a more abundant mass spectral feature. The left column of molecular structures (**A, C, E, G**) relate to *m/z* values of monophosphoribonucleotides (*i.e.,* RNA), while the right column (**B, D, F, H**) relates to monophosphodeoxyribonucleotides (*i.e.,* DNA). It should be noted that dehydration of the sugar moiety (*i.e.*, five-membered ribose or deoxyribose rings in red boxes) leads to the formation of a new carbon-carbon double bond, and thus nitrogen bases originating from nuclear (deoxyribose), or ribosomal (ribose) compartments, can be differentiated in the mass spectrum (*i.e.*, structural differences pre- and post-dehydration observed in the red boxes).

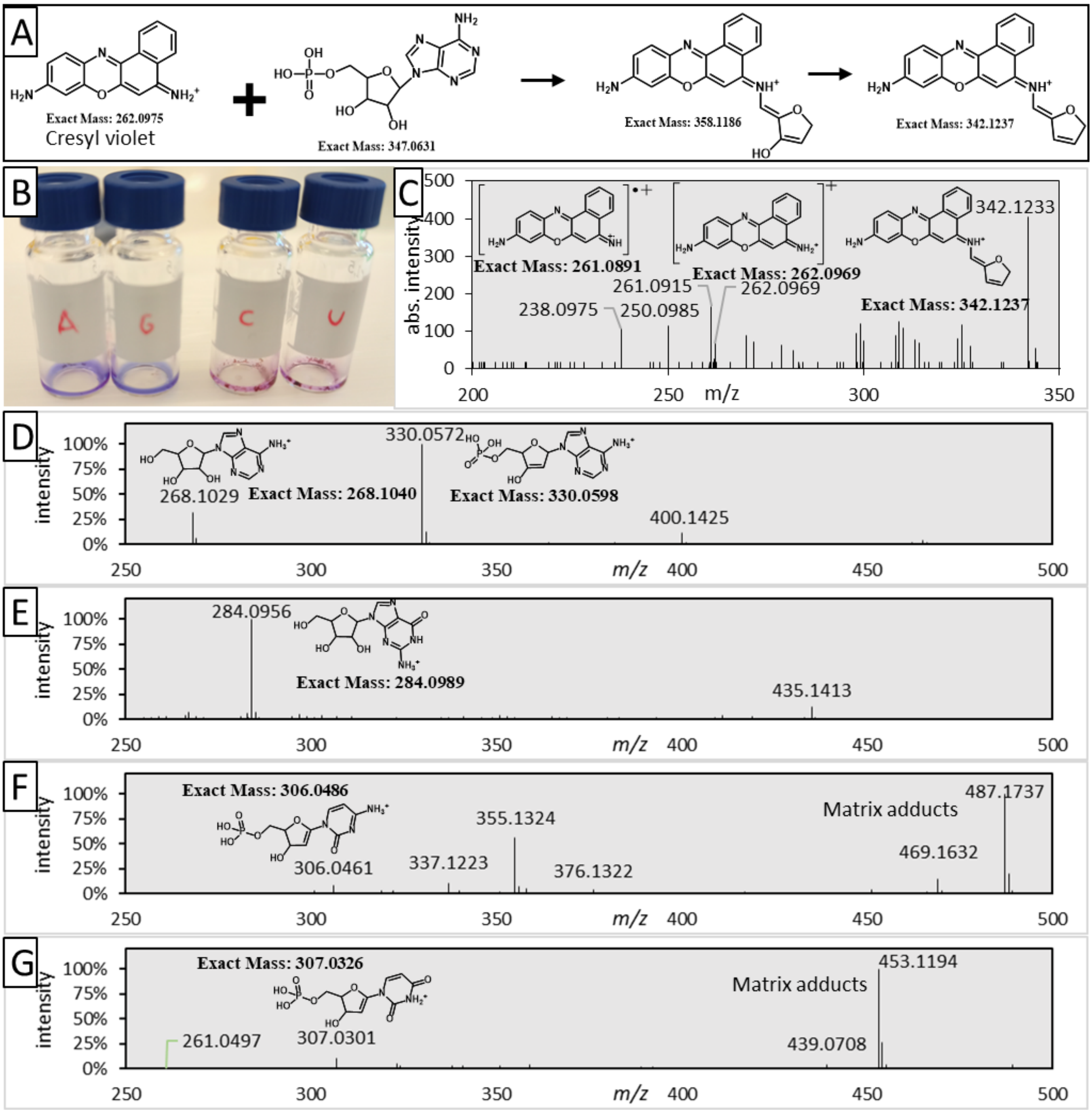

**Supplementary Fig. 6: CV derivative MS$^2$ and nucleotide standards AP-MALDI-P MS$^1$**

**(A)** Graphical representation of the formation of the *m/z* 342 mass spectral feature from cresyl violet and purine-based ribonucleotides. NB: the amount of conjugation is extended in the *m/z* 342 structure compared to the cresyl violet molecule (*m/z* 262). **(B)** Photograph of mixed cresyl violet with individual ribonucleotide standards. Adenine (A) and guanine (G) are classified as purines, and cytosine (C) and uracil (U) are pyrimidines. Notably, a colour change to deep blue/purple can be observed for the purines mixed with cresyl violet, while the pyrimidines mixed with cresyl violet are observed as a red/pink colour. This colour change to deep blue/purple with the purines and cresyl violet potentially indicates that there has been a change to the molecular conjugation and thus UV-Vis absorbance (*i.e.,* colour). **(C)** Tandem MS of the *m/z* 342 feature from murine spinal cord tissue displaying that the CV molecular ion (*m/z* 262) is one of the product ions. Additionally, a feature 1 Da less than CV appears as an abundant product ion (*m/z* 261). We speculate that interactions between the conjugated amines and the cold-plasma could generate a radical cation, that subsequently leads to ring-opening fragmentation observed as product ion *m/z* 238 and 250. **(D-G)** MS$^1$ spectra from spotted nucleotide standards without the addition of cresyl violet with spectral features corresponding to the molecular structure indicated. For adenine (D), guanine (E), cytosine (F) and uracil (G).

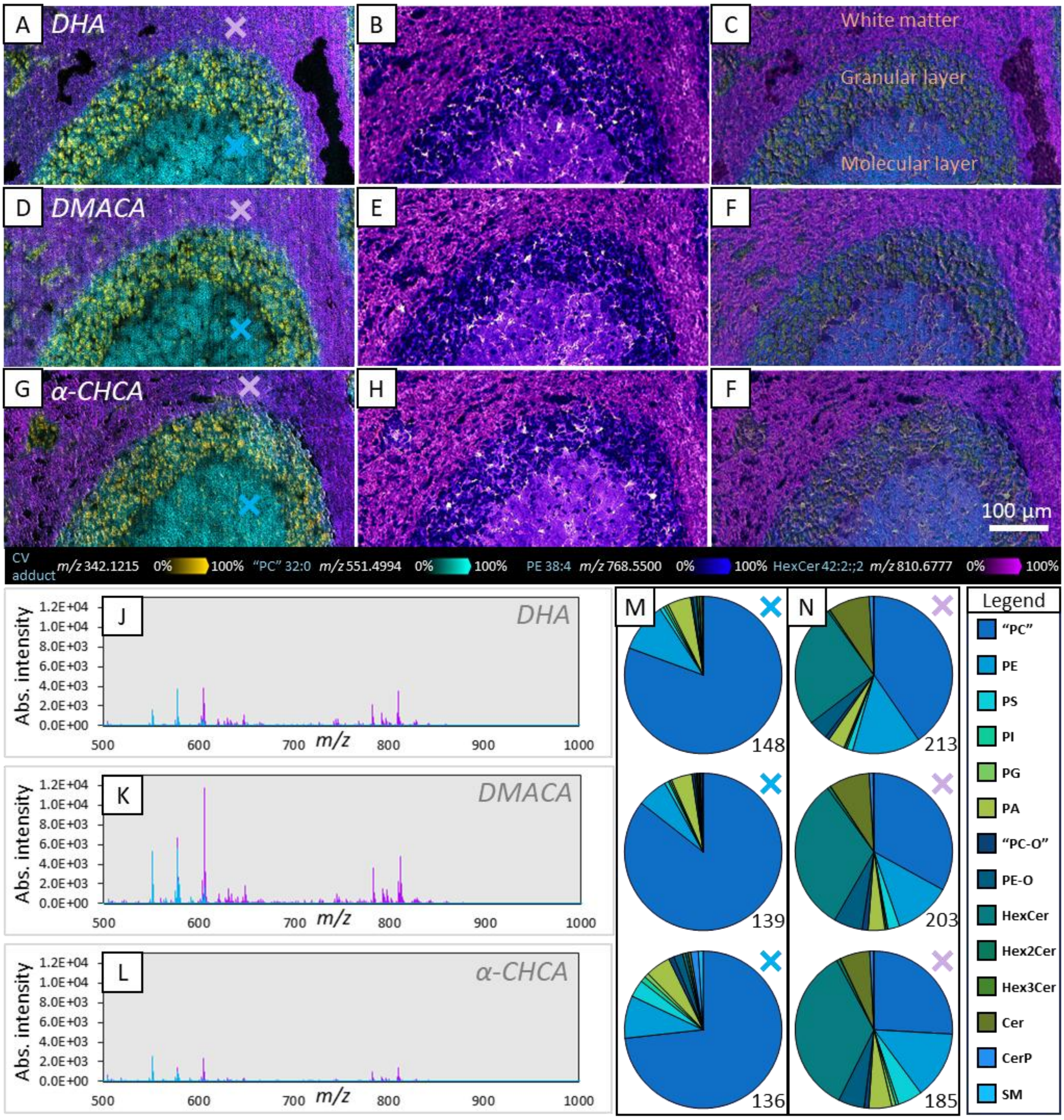

**Supplementary Fig. 7: Effect of matrix choice on MSI spatial resolution and lipid coverage within sagittal mouse cerebellar sections.**

**(A-C)** Representative ion images for 1 µm pixel size MSI using DHA matrix with CV pre-staining, 10× brightfield microscopy and overlay of both imaging modalities. **(D-F)** Representative ion images for 1 µm pixel size MSI using DMACA with CV pre-staining, and 10× brightfield microscopy; and composite of both imaging modalities. **(G-I)** Representative ion images for 1 µm pixel size MSI $\alpha$-CHCA matrix with CV pre-staining, 10× brightfield microscopy and overlay of both imaging modalities. In each MSI example the same *m/z* values are used for visualisation and provided in the key below. **(J-L)** Absolute intensity, averaged (5×5 pixels) mass spectra from similar tissue regions (indicated by purple and blue 'X') from the DHA (A), DMACA (D), and $\alpha$-CHCA (G) MS images. **(M)** Graphical comparison of total lipid identified in the blue 'X' pixel regions (cerebellar molecular layer). **(N)** Graphical comparison of total lipid identified in the purple 'X' pixel regions (cerebellar white matter). Lipid classes being monitored in the charts are annotated in the 'legend', and lipid species counts are annotated beneath the charts. Laser energy for each selected matrix was optimised for maximum sample ablation of the 1 µm pixel, without (or with very minimal) oversampling. All lipid identifications are assigned as $[M+H]^+$ ions, with the exception of "PCs", which are assigned as $[M-183.066+H]^+$ ions.

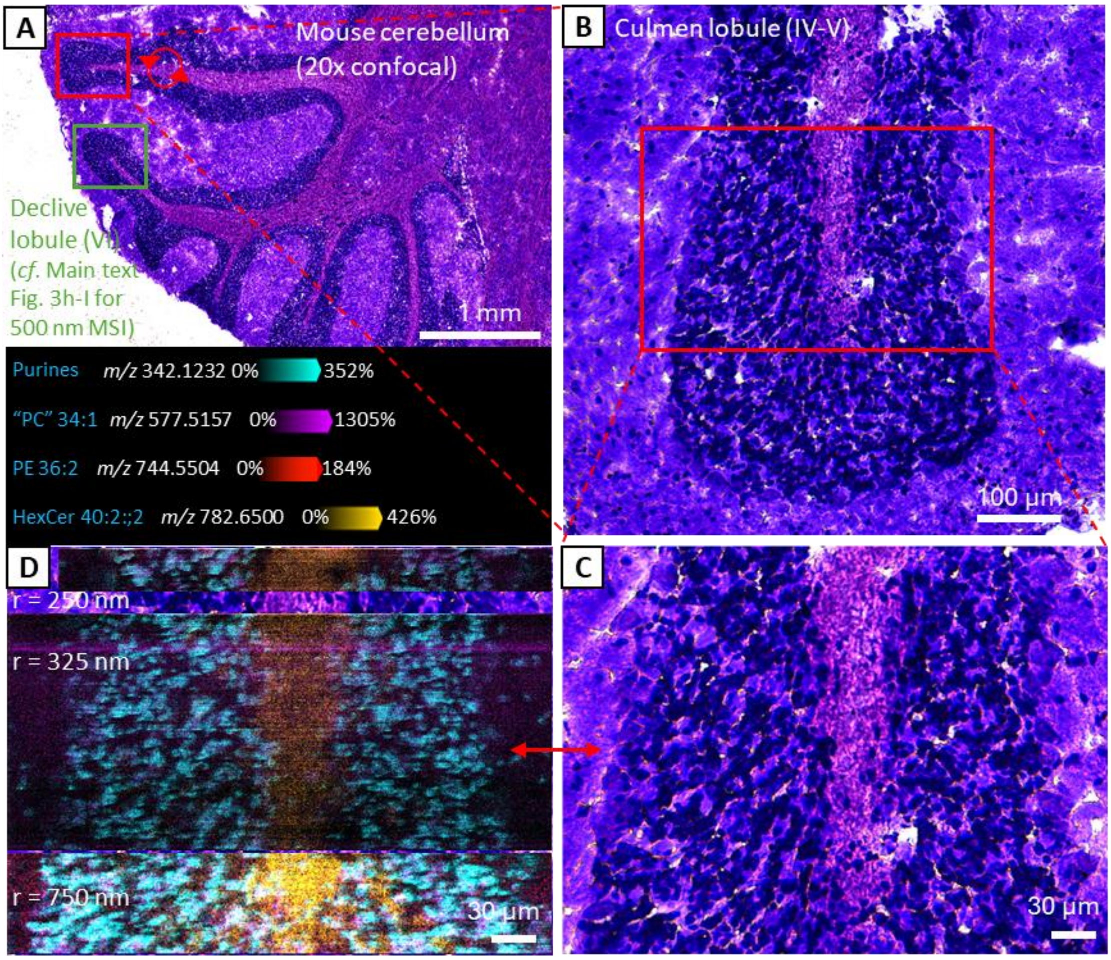

**Supplementary Fig. 8: Spatial resolution capabilities of the t-MALDI-P (750 nm, 500 nm, 325 nm 250 nm pixel sizes)**

**(A)** 20× confocal microscopy of a pre-stained sagittal section of mouse cerebellum showing the Culmen lobule (IV-V) in the red box, used throughout this figure, and the Declive lobule (green box) used for 500 nm MSI within Main text Fig. 3h-i. **(B)** Magnification of the Culmen lobule, with the red box indicating the region imaged at different MSI lateral resolutions (C-D). **(C)** 20× confocal microscopy of the tissue used for MSI. **(D)** from top to bottom: MSI images obtained using different pixel sizes of 250 nm, 325 nm and 750 nm. The key displayed the ions used to generate the images. While analyte ions are still visible in the 250 and 325 nm pixel size MS images, a significant drop in absolute intensity can be noted as these images are obtained using extreme oversampling conditions (*i.e.,* lateral resolution movement >50% of the nominal laser spot size). All lipid identifications are assigned as $[M+H]^+$ ions, with the exception of "PCs", which are assigned as $[M-183.066+H]^+$ ions.

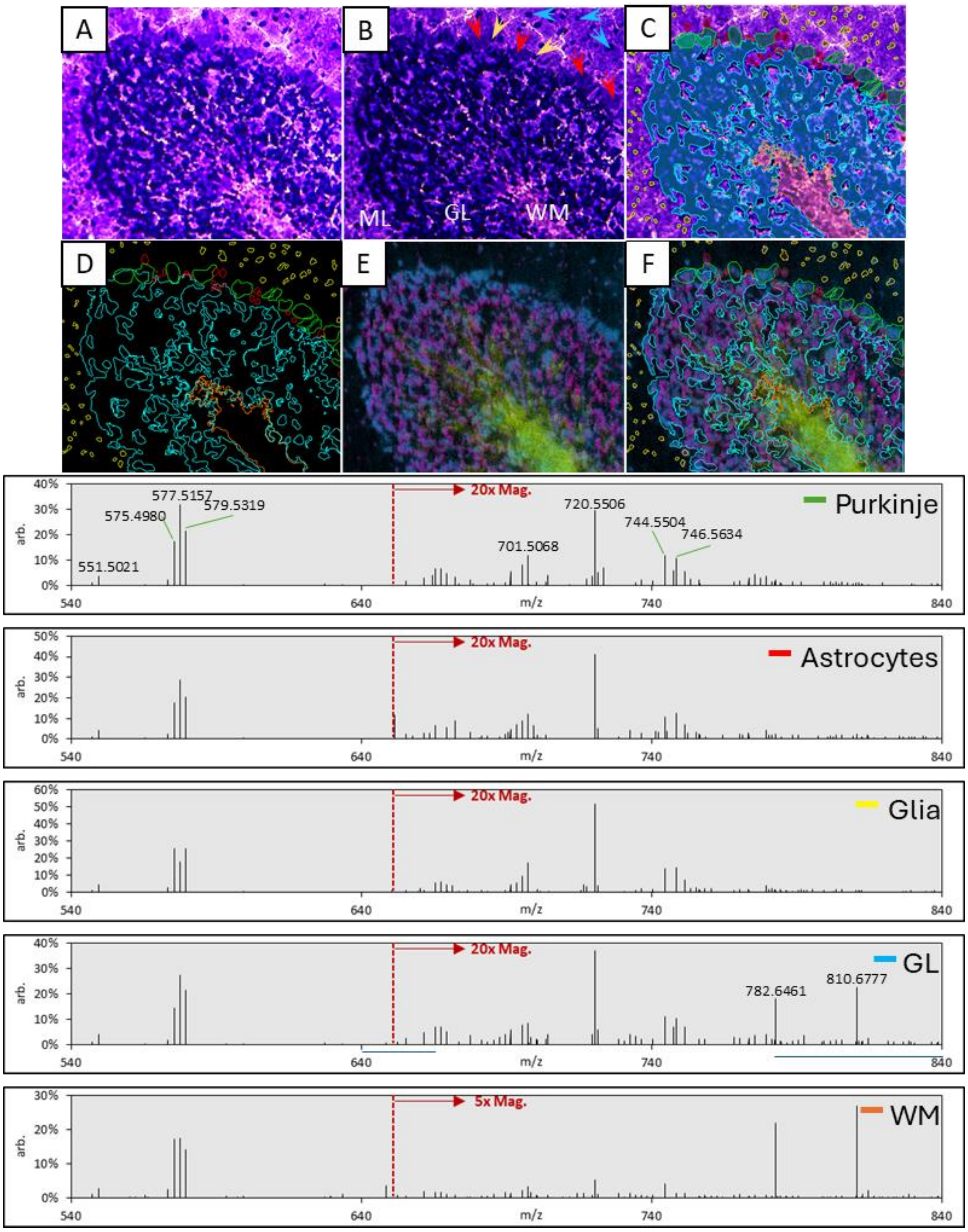

**Supplementary Fig. 9: QuPath region segmentation and averaged region mass spectra**

**(A)** 20x confocal microscopy image of CV pre-stained mouse cerebellum is obtained and imported into QuPath. **(B)** Cell types can then be identified within the optical images, such as the Purkinje neurons (red arrows), the Bergmann astrocytes (yellow arrows) and general glial cells of the outer molecular layer (blue arrows). **(C)** Regions of interest (ROI) can then be generated in QuPath based on the microscopy and user set identification parameters or manual ROI selection tools. **(D)** ROIs can then be saved and exported into MSI analytical software, such as SCiLS. **(E)** Using the microscopy image and an MSI image, a multi-point image co-registration can be undertaken. **(F)** Microscopy derived ROIs can then accurately segment MSI data for mass spectral investigation. Background subtracted averages from each of the colour-coded ROIs can be seen in the mass spectra above. GL = granular layer; WM = white matter.

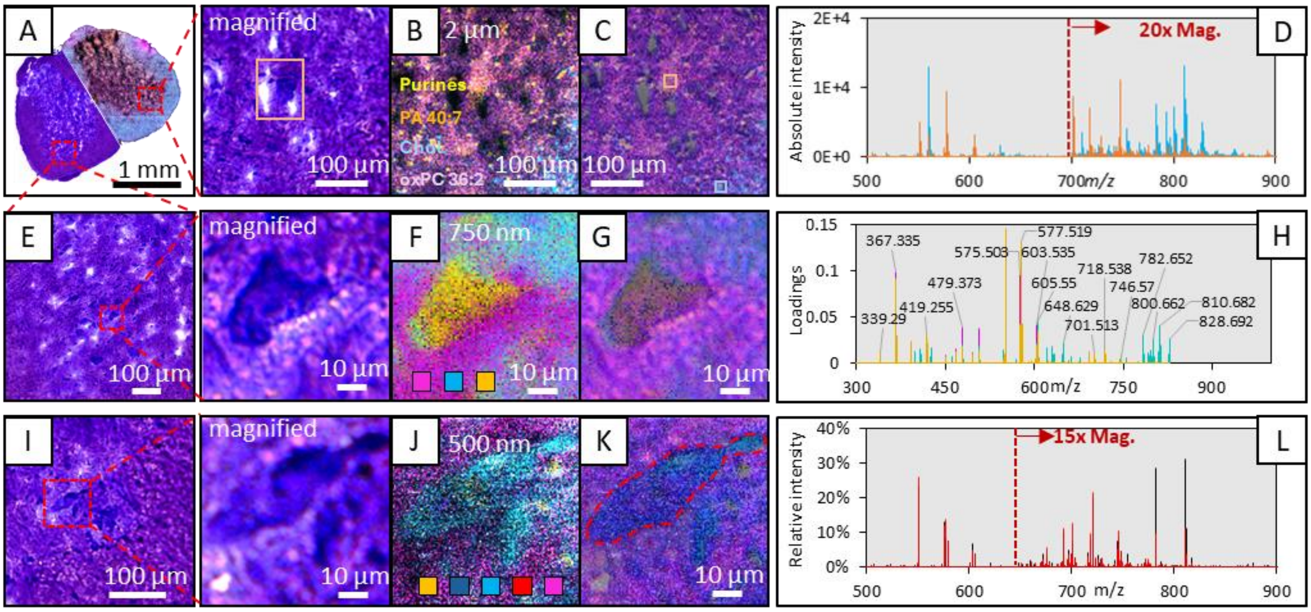

**Supplementary Fig. 10: Murine spinal cord and in situ motor neuron MSI using non-oversampling and oversampling conditions.**

**(A-C)** DMACA coated, cresyl violet pre-stained transverse murine spinal cord sections imaged through 10× brightfield microscopy (A) and MSI at 2 µm pixel size displaying purine nucleotides @ *m/z* 342.12 (yellow), PA 40:7 @ *m/z* 747.50 (orange), cholesterol @ *m/z* 369.35 (blue), and oxidised "PC" 36:2 @ *m/z* 493.39 (purple) (B). A composite image of both is displayed in (C). **(D)** Absolute intensity, averaged mass spectrum of 5×5-pixel regions indicated by the blue and orange boxes in (C). **(E-G)** *in situ* motor neuron cell located within the murine spinal cord section, imaged through 10× brightfield microscopy (E) and MSI at 750 nm (F), with a composite image displayed in (G). **(H)** Probabilistic latent semantic analysis (pLSA) loadings mass spectrum displays 3 components of a 5 component pLSA analysis, with the loadings spectra being colour matched to each component (yellow displays lipids that differentiate the neuron from the surrounding stroma [magenta] or other cell types [cyan]). **(I-J)** *in situ* motor neuron cell located within the murine spinal cord section, imaged through 10× brightfield microscopy (I) and MSI at 500 nm (J), with a composite image and motor neuron perimeter displayed in (K). Colour overlays in the MSI (J) indicate the spatial distribution of purines @ *m/z* 342.125 (yellow), cholesterol @ *m/z* 369.353 (navy blue) "PC" 34:1 @ *m/z* 577.519, (cyan) HexCer 40:2;2 @ *m/z* 782.651 (red), and HexCer 42:1;3 @ *m/z* 828.690 (magenta). **(L)** The overlaid mass spectra display the relative intensity spectral profile from within the cell region (red) compared to the remaining image region (black). All lipid identifications are assigned as $[M+H]^+$ ions, with the exception of "PC" and cholesterol, which are assigned as $[M-183.066+H]^+$ and $[M-H_2O+H]^+$, respectively.

**(Extended text for Supplementary Fig. 10). In situ single cell analysis capabilities.**

To demonstrate the capabilities of t-MALDI-P in analysing single cells found within tissues, fresh frozen mouse lumbar spinal cord sections were obtained. After pre-staining and recording microscopy, a hemisphere of a section was imaged at 2 µm (Supplementary Fig. 10A). Notably, within the micrographs, motor neurons (MN) are easily identified in the ventral horn as they are readily stained to black-purple by the cresyl violet dye and are >20 µm in size (as seen within the orange box in Supplementary Fig. 10A magnification). From the MSI (Supplementary Fig. 10B), numerous lipids and other uncharacterised ions could be identified, with the discrete signals being characteristic to tissue and cell types. For example, the putatively assigned PA 40:7 (orange; m/z 747.498) is more abundant in the MN, the oxidised (*i.e.*, oxidative cleavage of the carbon-carbon double bond in the fatty acids) "PC" 36:2 (pink; m/z 493.39) is largely found within the grey-matter stroma, and globular cholesterol (blue; m/z 369.353) features can be found in the white matter surrounding the ventral horn. Taking an average of the 5×5-pixel regions from within the single MN and white matter (Supplementary Fig. 10C; orange and blue boxes, respectively), the underlying mass spectra are compared in Supplementary Fig. 10D. High spectral intensity of glycosphingolipids can be observed in the white matter, while the MN has a greater spectral intensity of "PC", PE, and PA glycerophospholipids. It should be noted that PA assignments are likely ambiguous and could also partially arise from source fragmentation of lipids, such as PS. However, comparison of the ion distributions of intact PS lipids across the spinal cords tissue and the analogous PA lipids that would arise from PS fragmentation, reveals different spatial distributions of the two analyte assignments. This suggests that, at least in part, PA is not solely arising from PS fragmentation.

The *in situ* single cell imaging experiments were repeated on separate MNs located in the ventral horn at 750 nm and 500 nm lateral resolutions, with the multimodal images and spectra for each being displayed in Supplementary Fig. 10E-H and I-L, respectively. From the 750 nm MSI (Supplementary Fig. 10F) data, a feature list of approximately 400 unique *m/z* values (*i.e.*, spectral features, not lipid IDs) was generated using a spatial correlation (Pearson's; $P \leq 0.3$) to the highest on-cell signal (*i.e., m/z* 577.513). The feature list was subjected to a 5 component pLSA analysis, which identifies within the data (*i.e.,* mass spectral) commonality between pixels, and reports back weighted loadings as mass spectra. Three of these five components were used to generate the 750 nm MSI image (Supplementary Fig. 10F), with each components characterising unique tissue and cell types (*i.e.,* yellow displays lipids that differentiate the MN from the surrounding stroma [magenta] or other cell types [cyan]). From the 3 components displayed, a total of 96 spatially informative ions are observed within the loadings spectra (Supplementary Fig. 10H).

For Supplementary Fig. 10J, oversampling conditions were used to obtain MSI of a MN at 500 nm (*cf.* Supplementary Fig. 8 for cerebellum imaged at 250 nm also using oversampling). Here, total ion count normalisation was used to improve the visualisation of the 5 selected mass spectral features and their spatial distribution across the MSI region: $[\text{purines+H}]^+$ @ *m/z* 342.125 (yellow), $[\text{cholesterol-H}_2\text{O+H}]^+$ @ *m/z* 369.353 (navy blue), $[\text{“PC” 34:1 -183+H}]^+$ @ *m/z* 577.519 (cyan), $[\text{HexCer 40:2;2+H}]^+$ @ *m/z* 782.651 (red), and $[\text{HexCer 42:1;3 +H}]^+$ @ *m/z* 828.690 (magenta). Notably, at this pixel size distinct variation in lipid distribution could be observed across the MN cell, as is observed with the cholesterol (navy) and "PC" (cyan) lipids. Using the microscopy image (Supplementary Fig. 10I) to define the cell perimeter (Supplementary Fig. 10K) in the MSI (Supplementary Fig. 10J), the averaged mass spectra from the 3850 pixels region was generated (Supplementary Fig. 10L; red). For comparison, the average spectrum of the remaining tissue region was generated (Supplementary Fig. 10L; black). More than 27 lipid features could be identified within the whole tissue region (black), many of which are also found within the MN cell (red) but at differing intensities. Of the 27 lipids however, five ceramides and hexosylceramides were almost entirely absent from within the MN cell region compared to the surrounding tissue. While lower spatial resolution MSI experiments may have difficulty deconvoluting whether these lipids are also present around the perimeter of the cell or solely arising from the surrounding tissue, at 500 nm pixel sizes, this is easily differentiated.

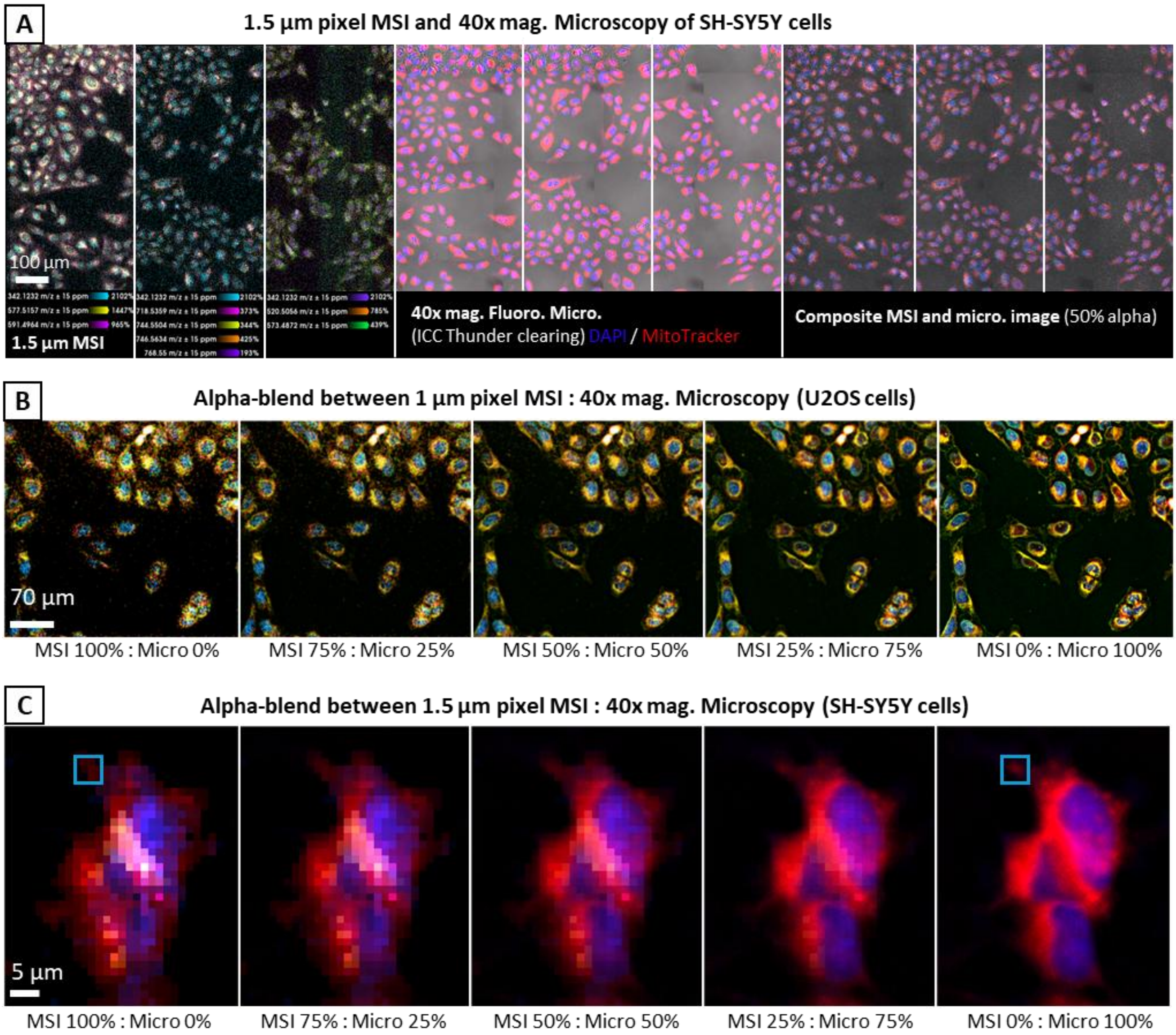

**Supplementary Fig. 11: Single cell co-registration and fidelity between MSI and 40x microscopy**

**(A)** Co-registration fidelity between 1.5 µm MSI and 40× fluorescence microscopy of SH-SY5Y neuroblastoma cells. Nine unique *m/z* lipid and nucleotide ion-images were selected for display in the MSI (left; *m/z* 342.123, 577.516, 591.496 , 718.536, 744.550, 746.63, 768.550, 520.506 and 573.487, *i.e.,* purine nucleotides, "PC" 34:1, "PC-O" 36:1, PE 34:1, PE 36:2, PE 36:1, PE 38:4, Cer 34:2;1, and "PC" 34:3, respectively). Microscopy (mid) displays the brightfield and 2-channel fluorescence microscopy of the cells, which had been stained with DAPI and MitoTracker Deep Red to reveal the location of cell nuclei and mitochondria, respectively. A composite of the two images, each with an alpha/transparency of 50%, is found on the right. **(B)** To assist visualisation of imaging co-registration fidelity, the 1 µm MSI and 40× fluorescence microscopy of U2OS osteosarcoma cells from Main text Fig. 4 is displayed with varying degrees of transparency (indicated below each image in the series). U2OS cells were stained with Hoechst and Nile red, which is visible in the blue, red/orange, and green/yellow channels in the microscopy. Four spectral features were selected within the MSI (*m/z* 342.123, 575.503, 577.519, and 603.535, *i.e.,* purine nucleotides, "PC" 34:2, "PC" 34:1, and "PC" 36:2, respectively) that were detected in the same regions as the fluorescence stains in the fluorescence microscopy. **(C)** An additional visualisation of imaging co-registration fidelity, this time showing varied transparency images of a magnified region across a group of SH-SY5Y cells to highlight the resolution capabilities of the t-MALDI-P at 1.5 µm pixel size compared to 40× fluorescence microscopy. As in (A), the SH-SY5Y cells were stained with DAPI and MitoTracker Deep Red, and the MSI shows 6 representative lipids and nucleotides (purine nucleotides, "PC" 34:2, PE 34:1, PE 34:0, PE 36:2, PE 36:1) that detected in the same regions as the fluorescent stains. All lipid identifications are assigned as $[M+H]^+$ ions, with the exception of "PCs", which are assigned as $[M-183.066+H]^+$.

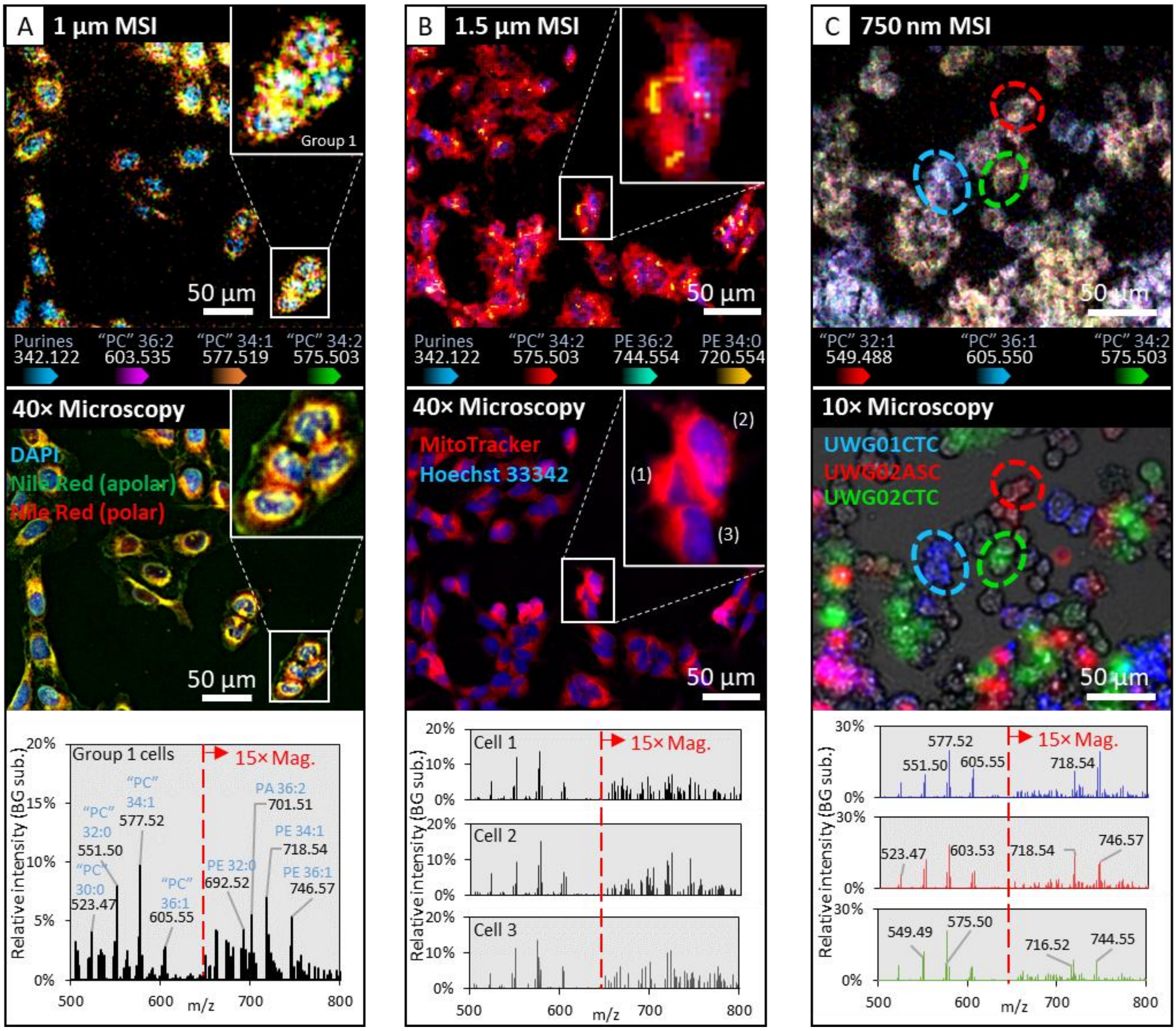

**Supplementary Fig. 12: Mass spectral investigation of single cells.**

**Mass spectrometry imaging of single cell cultures at near sub-cellular spatial resolutions. (A)** Osteosarcoma cell line (U2OS) stained with DAPI nuclear stain and the lipid stain, Nile red, and imaged through: MSI at 1 µm (top) and 3-channel (390/470 nm, 555/550 nm, 475/510 nm) 40× fluorescence microscopy (mid). The mass spectrum (bottom) displays the spectral features arising from "on-cell" regions in the "Group 1" cells determined through co-registration of the two imaging modalities. **(B)** Neuroblastoma cell line (SH-SY5Y) stained with the nuclear stain Hoechst 33342 and mitochondrial stain MitoTracker Deep Red and imaged through: MSI at 1.5 µm pixel size (top) and 2-channel (390/470 nm, 635/700 nm) 40× fluorescence microscopy (mid). The mass spectrum (bottom) displays the spectral features arising from "on-cell" regions determined through co-registration of the two imaging modalities. **(C)** Established patient derived circulating tumour cell lines originally isolated from peripheral venous blood (CTC) or ascites (ASC). JI lines (red/green) are from the same donor. Monocultures of each line were stained with different fluorescent CellBrite membrane dyes (red, green, blue) before being trypsinized and co-cultured (72hrs). Top-to-bottom: MSI at 750 nm pixel size, 3-channel (390/470 nm, 555/550 nm, 475/510 nm) 10× fluorescence microscopy and a composite mass spectrum displaying the colour-matched mass spectral features arising from the cells indicated in the red, green or blue ellipses. Red/green overlayed ellipses in the MSI indicate that cells were not able to be differentiated solely through their mass spectral profile.

Lipid identifications were based on $MS^1$ accurate mass ($\Delta ppm \leq 6$), with colour intensities displaying relative intensity with hotspot removal above the $99^{th}$ percentile. A 15-25× magnification was used to highlight lower-intensity spectral features consistent with PE and PE-O lipids in the m/z 650-800 mass range and the ordinate axis units are arbitrary due to background subtraction being used to simplify MS visualisation. All lipid identifications are assigned as $[M+H]^+$ ions, with the exception of "PCs", which are assigned as $[M-183.066+H]^+$.

### Supplementary tables

**Supplementary Table 1: Putatively assigned lipids identified within the 2 µm MSI of a conventionally prepared hippocampus section (cf. Main text Fig. 2c).**

IDs are based on accurate mass matching at the $MS^1$ level and thus some misassignments may be present. The ions nonetheless represent sample-related mass features and further support the rich biochemical information obtained.

PC assignments are based off the headgroup loss fragments (-183.066 *m/z*), and SM assignments are based off the loss of a methyl group (-14.016 *m/z*). Thus, all lipid identifications are assigned as $[M+H]^+$ ions, with the exception of PC, SM and cholesterol, which are assigned as $[M-183.066+H]^+$, $[M-14.016+H]^+$and $[M-H_2O+H]^+$, respectively. See the *Methods* for further details regarding source fragmentation of choline containing lipids.

NB: All lipid identifications are assigned from $[M+H]^+$ ions, with the exception of PC, SM and cholesterol, which are assigned from $[M-183.066+H]^+$, $[M-14.016+H]^+$and $[M-H_2O+H]^+$, respectively.

| Hippocampus no stain; 2 µm (164 IDs) | | | |
|---|---|---|---|
| Lipid ID | exp. m/z | Averaged abs. intensity (across whole area) | Δppm |
| PC 28:1 | 493.4225 | 13.7 | 5.43 |
| PC 30:1 | 521.4538 | 11.3 | 5.04 |
| PC 32:2 | 547.4689 | 15.9 | 5.79 |
| PC 32:1 | 549.4846 | 123.8 | 5.77 |
| PC 34:4 | 571.4692 | 2.6 | 4.97 |
| PC 34:3 | 573.4844 | 9.0 | 5.88 |
| PC 34:2 | 575.5008 | 161.0 | 4.44 |
| PC 34:1 | 577.5157 | 2273.3 | 5.73 |
| PC 36:4 | 599.4998 | 81.4 | 5.96 |
| PC 40:6 | 651.5314 | 123.1 | 5.00 |
| PC 40:2 | 659.5947 | 4.5 | 3.85 |
| PC 40:1 | 661.6096 | 5.2 | 5.07 |
| PC 42:6 | 679.5635 | 1.5 | 3.72 |
| PC 42:5 | 681.5779 | 2.0 | 5.41 |
| PC 42:4 | 683.5984 | 2.2 | 1.64 |
| PC 42:2 | 687.6264 | 2.7 | 3.23 |
| PC 42:1 | 689.6406 | 3.0 | 5.25 |
| PC 44:5 | 709.6101 | 1.2 | 4.05 |
| PC 44:4 | 711.6282 | 0.7 | 0.52 |
| PC 44:2 | 715.6603 | 0.4 | 0.57 |
| PE 28:2 | 632.4307 | 3.8 | 3.34 |
| PE 28:1 | 634.4450 | 2.8 | 1.25 |
| PE 30:2 | 660.4561 | 2.6 | 5.75 |
| PE 30:1 | 662.4735 | 2.0 | 3.00 |
| PE 32:3 | 686.4756 | 2.0 | 0.03 |
| PE 32:2 | 688.4899 | 2.2 | 1.89 |
| PE 32:1 | 690.5032 | 3.1 | 5.25 |
| PE 32:0 | 692.5190 | 12.4 | 5.09 |
| PE 34:6 | 708.4579 | 3.2 | 2.83 |
| PE 34:4 | 712.4934 | 3.0 | 3.07 |
| PE 34:3 | 714.5054 | 4.0 | 1.95 |

| | | | |
|---|---|---|---|
| PE 34:2 | 716.5196 | 6.9 | 4.02 |
| PE 34:1 | 718.5359 | 38.0 | 3.17 |
| PE 34:0 | 720.5506 | 203.8 | 4.46 |
| PE 36:5 | 738.5070 | 2.2 | 0.24 |
| PE 36:4 | 740.5185 | 8.9 | 5.40 |
| PE 36:3 | 742.5354 | 4.5 | 3.63 |
| PE 36:2 | 744.5504 | 31.2 | 4.50 |
| PE 38:7 | 762.5023 | 2.9 | 5.88 |
| PE 38:6 | 764.5180 | 18.9 | 5.85 |
| PE 38:4 | 768.5500 | 74.3 | 4.96 |
| PE 40:7 | 790.5355 | 7.2 | 3.32 |
| PE 40:6 | 792.5500 | 62.6 | 4.79 |
| PE 40:4 | 796.5824 | 12.8 | 3.37 |
| PE 40:1 | 802.6274 | 1.6 | 5.80 |
| PE 42:7 | 818.5673 | 4.5 | 2.66 |
| PE 42:5 | 822.5963 | 2.4 | 5.38 |
| PE 42:4 | 824.6183 | 1.7 | 2.28 |
| PE 42:3 | 826.6328 | 3.0 | 0.91 |
| PE 42:2 | 828.6481 | 3.7 | 0.49 |
| PE 42:1 | 830.6642 | 1.9 | 0.98 |
| PS 30:1 | 706.4628 | 3.9 | 3.56 |
| PS 32:2 | 732.4833 | 3.4 | 3.18 |
| PS 32:1 | 734.4968 | 3.2 | 0.15 |
| PS 34:6 | 752.4470 | 1.9 | 3.58 |
| PS 34:5 | 754.4625 | 3.0 | 3.78 |
| PS 34:3 | 758.4983 | 1.8 | 2.19 |
| PS 36:6 | 780.4772 | 2.3 | 4.83 |
| PS 36:5 | 782.4935 | 4.0 | 4.08 |
| PS 40:5 | 838.5553 | 1.1 | 4.70 |
| PS 40:4 | 840.5703 | 1.7 | 5.52 |
| PS 40:3 | 842.5901 | 0.9 | 0.59 |
| PS 40:2 | 844.6105 | 1.5 | 5.06 |
| PS 40:1 | 846.6231 | 2.5 | 1.40 |
| PS 40:0 | 848.6404 | 3.9 | 3.42 |
| PS 42:3 | 870.6243 | 1.9 | 2.81 |
| PS 42:2 | 872.6421 | 3.9 | 5.30 |
| PS 42:1 | 874.6559 | 5.2 | 3.12 |
| PS 42:0 | 876.6699 | 4.1 | 1.26 |
| PS 44:12 | 880.5138 | 1.0 | 1.63 |
| PS 44:6 | 892.6018 | 5.4 | 4.97 |
| PS 44:5 | 894.6169 | 2.6 | 5.59 |
| PS 44:3 | 898.6517 | 0.5 | 1.62 |
| PS 44:2 | 900.6714 | 0.5 | 2.93 |
| PS 44:1 | 902.6867 | 0.5 | 2.48 |
| PI 32:3 | 805.4818 | 1.0 | 5.45 |
| PI 36:1 | 865.5763 | 1.0 | 4.32 |

| | | | |
|---|---|---|---|
| PI 36:0 | 867.5911 | 3.6 | 5.28 |
| PI 38:8 | 879.5017 | 0.7 | 0.08 |
| PI 38:7 | 881.5137 | 0.8 | 4.26 |
| PI 38:1 | 893.6065 | 2.6 | 5.42 |
| PI 40:8 | 907.5336 | 0.5 | 0.49 |
| PI 42:10 | 931.5366 | 0.3 | 3.79 |
| PI 42:9 | 933.5463 | 0.3 | 2.67 |
| PI 42:4 | 943.6219 | 0.4 | 5.45 |
| PI 44:11 | 957.5525 | 0.2 | 3.88 |
| PI 44:10 | 959.5654 | 0.2 | 1.08 |
| PI 44:1 | 977.7026 | 0.2 | 2.74 |
| PG 42:10 | 843.5132 | 0.7 | 4.57 |
| PG 44:10 | 871.5477 | 0.6 | 0.81 |
| PA 34:6 | 665.4148 | 3.6 | 4.31 |
| PA 34:5 | 667.4307 | 2.6 | 3.88 |
| PA 36:8 | 689.4200 | 2.5 | 3.31 |
| PA 36:7 | 691.4326 | 2.3 | 1.12 |
| PA 36:6 | 693.4510 | 2.7 | 2.94 |
| PA 38:8 | 717.4517 | 3.2 | 3.76 |
| PA 38:7 | 719.4634 | 2.7 | 1.77 |
| PA 38:6 | 721.4771 | 6.8 | 4.45 |
| PA 38:5 | 723.4928 | 17.4 | 4.32 |
| PA 38:3 | 727.5230 | 4.4 | 5.83 |
| PA 40:8 | 745.4780 | 4.0 | 3.09 |
| PA 40:7 | 747.4935 | 15.1 | 3.27 |
| PA 40:5 | 751.5258 | 23.9 | 1.86 |
| PA 42:9 | 771.4953 | 1.7 | 0.82 |
| PA 42:8 | 773.5115 | 2.9 | 0.06 |
| PA 42:7 | 775.5253 | 6.6 | 2.52 |
| PA 42:5 | 779.5607 | 6.4 | 2.80 |
| PA 44:11 | 795.4958 | 1.7 | 0.16 |
| PA 44:10 | 797.5110 | 1.9 | 0.79 |
| PA 44:9 | 799.5272 | 2.2 | 0.02 |
| PA 44:8 | 801.5446 | 1.4 | 2.10 |
| PA 44:7 | 803.5590 | 1.8 | 0.54 |
| PC-O 28:1 | 479.4433 | 4.6 | 5.28 |
| PC-O 28:0 | 481.4588 | 2.8 | 5.59 |
| PC-O 32:1 | 535.5079 | 4.0 | 1.02 |
| PC-O 36:2 | 589.5536 | 5.2 | 3.07 |
| PC-O 38:4 | 613.5524 | 2.9 | 4.94 |
| PC-O 38:3 | 615.5681 | 1.5 | 4.76 |
| PC-O 40:4 | 641.5843 | 1.8 | 3.72 |
| PC-O 42:7 | 663.5708 | 2.0 | 0.33 |
| PC-O 42:6 | 665.5878 | 2.4 | 1.69 |
| PC-O 42:5 | 667.6010 | 2.0 | 2.12 |
| PC-O 42:4 | 669.6202 | 1.4 | 3.19 |

| | | | |
|---|---|---|---|
| PC-O 44:7 | 691.6020 | 1.2 | 0.57 |
| PE-O 32:3 | 672.4956 | 2.2 | 0.94 |
| PE-O 34:5 | 696.4985 | 3.4 | 3.18 |
| PE-O 34:4 | 698.5143 | 12.0 | 3.37 |
| PE-O 36:5 | 724.5245 | 5.3 | 4.21 |
| PE-O 36:4 | 726.5451 | 3.8 | 2.63 |
| PE-O 36:3 | 728.5568 | 15.6 | 2.83 |
| PE-O 36:2 | 730.5704 | 15.1 | 5.64 |
| PE-O 38:5 | 752.5561 | 13.0 | 3.63 |
| PE-O 38:4 | 754.5719 | 5.2 | 3.45 |
| PE-O 40:4 | 782.6030 | 2.6 | 3.56 |
| PE-O 40:3 | 784.6169 | 1.6 | 5.82 |
| PE-O 40:2 | 786.6360 | 1.2 | 1.47 |
| PE-O 42:5 | 808.6212 | 1.7 | 0.34 |
| PE-O 42:4 | 810.6412 | 2.8 | 5.07 |
| PE-O 42:3 | 812.6541 | 3.4 | 1.65 |
| PE-O 42:2 | 814.6720 | 10.0 | 4.39 |
| PE-O 42:1 | 816.6867 | 2.5 | 3.24 |
| PE-O 44:4 | 838.6685 | 1.4 | 0.14 |
| PE-O 44:3 | 840.6880 | 1.3 | 4.63 |
| PE-O 44:2 | 842.7038 | 1.1 | 4.87 |
| HexCer 34:2;2 | 698.5527 | 1.2 | 5.51 |
| HexCer 36:2;2 | 726.5851 | 25.1 | 3.79 |
| HexCer 40:2;2 | 782.6461 | 196.6 | 5.59 |
| HexCer 42:2;2 | 810.6777 | 183.3 | 4.98 |
| Hex2Cer 28:0;2 | 780.5475 | 4.7 | 0.92 |
| Hex2Cer 30:2;2 | 804.5480 | 4.0 | 1.47 |
| Hex2Cer 30:1;2 | 806.5659 | 4.2 | 4.28 |
| Hex2Cer 30:0;2 | 808.5808 | 2.6 | 3.33 |
| Hex2Cer 32:2;2 | 832.5810 | 1.5 | 3.57 |
| Hex2Cer 32:1;2 | 834.5942 | 1.5 | 0.52 |
| Hex2Cer 34:2;2 | 860.6049 | 1.4 | 5.16 |
| Hex2Cer 34:1;2 | 862.6211 | 1.8 | 4.54 |
| Hex2Cer 34:0;2 | 864.6377 | 1.6 | 3.46 |
| Hex2Cer 40:2;2 | 944.7029 | 0.2 | 0.35 |
| Hex2Cer 40:0;2 | 948.7360 | 0.2 | 1.46 |
| Hex2Cer 42:2;2 | 972.7290 | 0.2 | 5.73 |

| | | | |
|---|---|---|---|
| Hex2Cer 42:1;2 | 974.7446 | 0.2 | 5.74 |
| Hex2Cer 44:2;2 | 1000.7655 | 0.2 | 0.35 |
| Hex2Cer 44:0;2 | 1004.7917 | 0.1 | 5.47 |
| Hex3Cer 30:1;2 | 968.6134 | 0.3 | 1.95 |
| Hex3Cer 32:2;2 | 994.6347 | 0.1 | 3.80 |
| Hex3Cer 32:1;2 | 996.6509 | 0.1 | 4.32 |
| Hex3Cer 32:0;2 | 998.6661 | 0.1 | 3.93 |
| Hex3Cer 36:2;2 | 1050.6880 | 0.1 | 5.20 |
| Hex3Cer 36:0;2 | 1054.7252 | 0.1 | 0.34 |
| Hex3Cer 38:1;2 | 1080.7392 | 0.1 | 1.20 |
| Hex3Cer 38:0;2 | 1082.7566 | 0.1 | 0.47 |
| Hex3Cer 42:1;2 | 1136.8033 | 0.0 | 0.21 |
| Hex3Cer 44:2;2 | 1162.8224 | 0.0 | 3.22 |
| Cer 28:2;2 | 452.4086 | 3.6 | 2.71 |
| Cer 32:2;2 | 508.4748 | 1.9 | 4.73 |
| Cer 36:2;2 | 564.5340 | 20.7 | 1.74 |
| Cer 36:1;2 | 566.5474 | 12.7 | 5.85 |
| Cer 42:2;2 | 648.6256 | 68.2 | 5.15 |
| CerP 28:2;2 | 532.3737 | 3.9 | 4.60 |
| CerP 30:2;2 | 560.4055 | 6.7 | 3.44 |
| CerP 30:1;2 | 562.4210 | 11.2 | 3.74 |
| CerP 32:2;2 | 588.4404 | 2.8 | 2.82 |
| CerP 32:1;2 | 590.4534 | 2.3 | 1.75 |
| CerP 34:2;2 | 616.4675 | 1.5 | 4.10 |
| CerP 36:1;2 | 646.5177 | 1.7 | 1.14 |
| CerP 38:2;2 | 672.5293 | 15.6 | 5.04 |
| CerP 40:2;2 | 700.5639 | 2.0 | 0.05 |
| CerP 40:1;2 | 702.5809 | 1.6 | 1.90 |
| CerP 42:2;2 | 728.5969 | 6.2 | 2.23 |
| CerP 42:1;2 | 730.6142 | 2.2 | 4.56 |
| CerP 44:2;2 | 756.6234 | 3.3 | 4.22 |
| CerP 44:1;2 | 758.6387 | 0.8 | 4.67 |
| SM 34:1;2 | 689.5579 | 1.7 | 1.94 |
| SM 36:2;2 | 715.5708 | 2.1 | 5.60 |
| SM 36:1;2 | 717.5880 | 16.2 | 3.47 |
| SM 38:1;2 | 745.6234 | 1.7 | 2.09 |

**Supplementary Table 2: Putatively assigned lipids identified within the 2 µm MSI of a hippocampus section having undergone novel pre-staining preparation (cf. Main text Fig. 2f).**

IDs are based on accurate mass matching at the $MS^1$ level and thus some misassignments may be present. The ions nonetheless represent sample-related mass features and further support the rich biochemical information obtained.

PC assignments are based off the headgroup loss fragments (-183.066 *m/z*), and SM assignments are based off the loss of a methyl group (-14.016 *m/z*). Thus, all lipid identifications are assigned as $[M+H]^+$ ions, with the exception of PC, SM and cholesterol, which are assigned as $[M-183.066+H]^+$, $[M-14.016+H]^+$and $[M-H_2O+H]^+$, respectively. See the *Methods* for further details regarding source fragmentation of choline containing lipids.

NB: All lipid identifications are assigned from $[M+H]^+$ ions, with the exception of PC, SM and cholesterol, which are assigned from $[M-183.066+H]^+$, $[M-14.016+H]^+$and $[M-H_2O+H]^+$, respectively.

| Hippocampus CV stain; 2 µm (213 IDs) | | | |
|---|---|---|---|
| Lipid ID | exp. m/z | Averaged abs. intensity (across whole area) | Δppm |
| PC 28:1 | 493.4225 | 193.5 | 5.43 |
| PC 30:1 | 521.4538 | 130.9 | 5.04 |
| PC 32:2 | 547.4689 | 278.0 | 5.79 |
| PC 32:1 | 549.4846 | 1259.1 | 5.77 |
| PC 34:6 | 567.4375 | 11.6 | 5.73 |
| PC 34:4 | 571.4692 | 39.0 | 4.97 |
| PC 34:3 | 573.4844 | 425.6 | 5.88 |
| PC 34:2 | 575.5008 | 4480.5 | 4.44 |
| PC 34:1 | 577.5157 | 13863.9 | 5.73 |
| PC 36:4 | 599.4998 | 590.9 | 5.96 |
| PC 38:7 | 621.4843 | 59.0 | 5.53 |
| PC 38:4 | 627.5326 | 847.7 | 3.30 |
| PC 38:3 | 629.5471 | 137.8 | 5.17 |
| PC 38:1 | 633.5796 | 224.6 | 3.21 |
| PC 40:6 | 651.5314 | 829.2 | 5.00 |
| PC 40:4 | 655.5638 | 113.3 | 3.33 |
| PC 40:2 | 659.5947 | 71.5 | 3.85 |
| PC 40:1 | 661.6096 | 84.5 | 5.07 |
| PC 42:6 | 679.5669 | 6.6 | 1.28 |
| PC 42:5 | 681.5779 | 11.0 | 5.41 |
| PC 42:4 | 683.5984 | 14.6 | 1.64 |
| PC 42:3 | 685.6111 | 15.3 | 2.62 |
| PC 42:2 | 687.6264 | 44.8 | 3.23 |
| PC 42:1 | 689.6406 | 56.4 | 5.25 |
| PC 44:6 | 707.5976 | 5.2 | 0.48 |
| PC 44:4 | 711.6282 | 3.2 | 0.52 |
| PC 44:3 | 713.6414 | 2.3 | 3.96 |
| PC 44:2 | 715.6567 | 3.1 | 4.43 |
| PC 44:0 | 719.6900 | 0.8 | 1.61 |
| PE 28:2 | 632.4307 | 19.7 | 3.34 |
| PE 28:1 | 634.4419 | 14.2 | 3.75 |

| | | | |
|---|---|---|---|
| PE 30:2 | 660.4561 | 14.6 | 5.75 |
| PE 30:1 | 662.4735 | 10.0 | 3.00 |
| PE 32:3 | 686.4756 | 12.9 | 0.03 |
| PE 32:1 | 690.5032 | 24.0 | 5.25 |
| PE 32:0 | 692.5190 | 124.2 | 5.09 |
| PE 34:6 | 708.4579 | 13.8 | 2.83 |
| PE 34:5 | 710.4728 | 12.1 | 3.90 |
| PE 34:4 | 712.4934 | 18.4 | 3.07 |
| PE 34:3 | 714.5054 | 31.2 | 1.95 |
| PE 34:2 | 716.5196 | 167.4 | 4.02 |
| PE 34:1 | 718.5359 | 281.0 | 3.17 |
| PE 34:0 | 720.5506 | 613.9 | 4.46 |
| PE 36:6 | 736.4900 | 9.9 | 1.67 |
| PE 36:5 | 738.5033 | 31.4 | 4.76 |
| PE 36:4 | 740.5185 | 93.2 | 5.40 |
| PE 36:3 | 742.5354 | 256.2 | 3.63 |
| PE 36:2 | 744.5504 | 581.6 | 4.50 |
| PE 36:1 | 746.5672 | 539.1 | 3.03 |
| PE 38:8 | 760.4882 | 10.9 | 3.93 |
| PE 38:7 | 762.5023 | 56.1 | 5.88 |
| PE 38:6 | 764.5180 | 197.4 | 5.85 |
| PE 38:5 | 766.5352 | 225.1 | 3.86 |
| PE 38:4 | 768.5500 | 632.6 | 4.96 |
| PE 38:3 | 770.5662 | 162.9 | 4.18 |
| PE 40:8 | 788.5183 | 32.4 | 5.34 |
| PE 40:7 | 790.5355 | 130.3 | 3.32 |
| PE 40:6 | 792.5500 | 490.2 | 4.79 |
| PE 40:4 | 796.5824 | 94.0 | 3.37 |
| PE 40:3 | 798.5963 | 27.2 | 5.54 |
| PE 40:2 | 800.6153 | 17.1 | 1.34 |
| PE 40:1 | 802.6274 | 7.1 | 5.80 |
| PE 42:5 | 822.5963 | 12.8 | 5.38 |
| PE 42:4 | 824.6183 | 9.8 | 2.28 |
| PE 42:3 | 826.6369 | 30.8 | 5.91 |
| PE 42:2 | 828.6481 | 39.2 | 0.49 |
| PE 42:1 | 830.6642 | 11.5 | 0.98 |
| PE 42:0 | 832.6810 | 2.9 | 2.37 |
| PE 44:4 | 852.6512 | 3.8 | 4.12 |
| PE 44:3 | 854.6658 | 4.8 | 2.92 |
| PS 30:2 | 704.4488 | 29.4 | 1.34 |
| PS 30:1 | 706.4664 | 21.8 | 1.44 |
| PS 32:3 | 730.4681 | 28.6 | 3.79 |
| PS 32:2 | 732.4833 | 29.7 | 3.18 |
| PS 32:1 | 734.5004 | 24.5 | 5.15 |
| PS 34:6 | 752.4508 | 16.3 | 1.42 |
| PS 34:5 | 754.4663 | 18.5 | 1.22 |

| | | | |
|---|---|---|---|
| PS 34:4 | 756.4834 | 14.0 | 3.16 |
| PS 34:3 | 758.4983 | 12.7 | 2.19 |
| PS 34:2 | 760.5148 | 10.3 | 3.29 |
| PS 36:6 | 780.4811 | 18.7 | 0.17 |
| PS 36:5 | 782.4935 | 25.8 | 4.08 |
| PS 36:4 | 784.5110 | 9.2 | 1.69 |
| PS 36:3 | 786.5258 | 8.3 | 2.70 |
| PS 38:4 | 812.5403 | 8.9 | 4.02 |
| PS 38:3 | 814.5579 | 7.3 | 1.62 |
| PS 40:6 | 836.5410 | 16.7 | 3.10 |
| PS 40:5 | 838.5553 | 8.8 | 4.70 |
| PS 40:4 | 840.5703 | 29.0 | 5.52 |
| PS 40:3 | 842.5901 | 6.2 | 0.59 |
| PS 40:2 | 844.6105 | 10.4 | 5.06 |
| PS 40:0 | 848.6404 | 52.2 | 3.42 |
| PS 42:3 | 870.6243 | 18.4 | 2.81 |
| PS 42:2 | 872.6421 | 53.2 | 5.30 |
| PS 42:1 | 874.6559 | 65.0 | 3.12 |
| PS 44:12 | 880.5138 | 6.9 | 1.63 |
| PS 44:7 | 890.5868 | 10.3 | 4.26 |
| PS 44:6 | 892.6018 | 53.6 | 4.97 |
| PS 44:5 | 894.6169 | 46.2 | 5.59 |
| PS 44:3 | 898.6517 | 2.8 | 1.62 |
| PS 44:2 | 900.6714 | 4.1 | 2.93 |
| PS 44:1 | 902.6867 | 2.4 | 2.48 |
| PI 36:1 | 865.5763 | 9.0 | 4.32 |
| PI 36:0 | 867.5911 | 41.0 | 5.28 |
| PI 38:8 | 879.5017 | 5.3 | 0.08 |
| PI 38:7 | 881.5181 | 6.3 | 0.74 |
| PI 38:6 | 883.5303 | 8.3 | 3.22 |
| PI 38:4 | 887.5595 | 23.3 | 5.55 |
| PI 38:1 | 893.6065 | 25.2 | 5.42 |
| PI 40:8 | 907.5336 | 3.5 | 0.49 |
| PI 42:0 | 951.6864 | 1.8 | 3.42 |
| PI 44:11 | 957.5525 | 1.6 | 3.88 |
| PI 44:1 | 977.7026 | 1.3 | 2.74 |
| PG 42:10 | 843.5174 | 4.2 | 0.43 |
| PG 44:10 | 871.5477 | 4.0 | 0.81 |
| PA 34:6 | 665.4148 | 19.7 | 4.31 |
| PA 34:5 | 667.4307 | 15.8 | 3.88 |
| PA 36:8 | 689.4200 | 13.9 | 3.31 |
| PA 36:7 | 691.4326 | 15.7 | 1.12 |
| PA 36:6 | 693.4476 | 19.3 | 2.06 |
| PA 36:5 | 695.4615 | 42.8 | 4.55 |
| PA 36:4 | 697.4778 | 83.4 | 3.63 |
| PA 36:1 | 703.5230 | 84.6 | 5.98 |

| | | | |
|---|---|---|---|
| PA 38:8 | 717.4481 | 23.8 | 1.24 |
| PA 38:6 | 721.4771 | 103.4 | 4.45 |
| PA 38:5 | 723.4928 | 240.2 | 4.32 |
| PA 40:8 | 745.4780 | 61.2 | 3.09 |
| PA 40:7 | 747.4935 | 223.3 | 3.27 |
| PA 40:4 | 753.5389 | 89.1 | 5.34 |
| PA 42:9 | 771.4953 | 11.7 | 0.82 |
| PA 42:8 | 773.5115 | 24.6 | 0.06 |
| PA 42:7 | 775.5253 | 42.5 | 2.52 |
| PA 42:6 | 777.5443 | 37.5 | 1.78 |
| PA 42:5 | 779.5607 | 26.0 | 2.80 |
| PA 42:4 | 781.5707 | 7.8 | 4.49 |
| PA 44:10 | 797.5110 | 15.0 | 0.79 |
| PA 44:9 | 799.5272 | 17.5 | 0.02 |
| PA 44:8 | 801.5446 | 14.6 | 2.10 |
| PA 44:7 | 803.5590 | 14.2 | 0.54 |
| PA 44:6 | 805.5744 | 13.9 | 0.28 |
| PC-O 34:5 | 555.4789 | 5.3 | 3.17 |
| PC-O 36:2 | 589.5536 | 21.1 | 3.07 |
| PC-O 38:4 | 613.5524 | 7.9 | 4.94 |
| PC-O 40:4 | 641.5843 | 3.8 | 3.72 |
| PC-O 42:5 | 667.6043 | 8.5 | 2.88 |
| PC-O 42:4 | 669.6202 | 4.8 | 3.19 |
| PC-O 42:3 | 671.6320 | 1.1 | 2.43 |
| PC-O 44:7 | 691.6020 | 4.5 | 0.57 |
| PC-O 44:6 | 693.6175 | 3.5 | 0.79 |
| PC-O 44:5 | 695.6354 | 2.2 | 2.43 |
| PC-O 44:4 | 697.6487 | 1.4 | 0.96 |
| PE-O 32:3 | 672.4956 | 13.9 | 0.94 |
| PE-O 34:3 | 700.5289 | 34.2 | 1.89 |
| PE-O 36:5 | 724.5318 | 17.8 | 5.79 |
| PE-O 36:4 | 726.5451 | 30.1 | 2.63 |
| PE-O 36:3 | 728.5568 | 55.6 | 2.83 |
| PE-O 36:2 | 730.5704 | 54.6 | 5.64 |
| PE-O 36:1 | 732.5859 | 19.1 | 5.83 |
| PE-O 38:5 | 752.5561 | 26.1 | 3.63 |
| PE-O 38:4 | 754.5719 | 22.8 | 3.45 |
| PE-O 40:5 | 780.5865 | 16.5 | 4.68 |
| PE-O 40:4 | 782.6069 | 11.5 | 1.44 |
| PE-O 40:3 | 784.6208 | 7.3 | 0.82 |
| PE-O 40:2 | 786.6399 | 8.2 | 3.53 |
| PE-O 42:5 | 808.6252 | 9.4 | 4.66 |
| PE-O 42:4 | 810.6412 | 20.7 | 5.07 |
| PE-O 42:2 | 814.6720 | 149.3 | 4.39 |
| PE-O 42:1 | 816.6867 | 25.0 | 3.24 |
| PE-O 44:4 | 838.6727 | 7.5 | 5.14 |

| | | | |
|---|---|---|---|
| PE-O 44:3 | 840.6880 | 9.6 | 4.63 |
| PE-O 44:2 | 842.7038 | 10.3 | 4.87 |
| PE-O 44:1 | 844.7161 | 1.8 | 0.83 |
| HexCer 34:2;2 | 698.5527 | 7.6 | 5.51 |
| HexCer 34:0;2 | 702.5845 | 11.2 | 4.83 |
| HexCer 36:2;2 | 726.5851 | 343.7 | 3.79 |
| HexCer 38:2;2 | 754.6172 | 199.6 | 2.59 |
| HexCer 40:2;2 | 782.6461 | 2095.2 | 5.59 |
| HexCer 42:2;2 | 810.6777 | 2293.5 | 4.98 |
| Hex2Cer 30:2;2 | 804.5480 | 8.5 | 1.47 |
| Hex2Cer 30:1;2 | 806.5659 | 12.1 | 4.28 |
| Hex2Cer 32:2;2 | 832.5769 | 6.5 | 1.43 |
| Hex2Cer 32:1;2 | 834.5983 | 7.6 | 5.52 |
| Hex2Cer 34:2;2 | 860.6049 | 7.9 | 5.16 |
| Hex2Cer 34:1;2 | 862.6211 | 15.7 | 4.54 |
| Hex2Cer 34:0;2 | 864.6377 | 15.9 | 3.46 |
| Hex2Cer 36:2;2 | 888.6385 | 23.0 | 2.42 |
| Hex2Cer 36:1;2 | 890.6536 | 51.4 | 3.09 |
| Hex2Cer 38:1;2 | 918.6825 | 6.9 | 5.62 |
| Hex2Cer 40:2;2 | 944.7029 | 1.2 | 0.35 |
| Hex2Cer 40:0;2 | 948.7360 | 0.7 | 1.46 |
| Hex2Cer 42:2;2 | 972.7290 | 1.3 | 5.73 |
| Hex2Cer 42:1;2 | 974.7446 | 1.1 | 5.74 |
| Hex2Cer 44:2;2 | 1000.7605 | 0.9 | 5.35 |
| Hex3Cer 30:2;2 | 966.6007 | 2.0 | 1.18 |
| Hex3Cer 30:1;2 | 968.6182 | 2.1 | 3.05 |
| Hex3Cer 32:2;2 | 994.6297 | 1.2 | 1.20 |
| Hex3Cer 36:0;2 | 1054.7252 | 0.5 | 0.34 |
| Hex3Cer 38:1;2 | 1080.7392 | 0.5 | 1.20 |
| Hex3Cer 38:0;2 | 1082.7512 | 0.5 | 4.53 |

| | | | |
|---|---|---|---|
| Cer 36:2;2 | 564.5340 | 146.0 | 1.74 |
| Cer 36:1;2 | 566.5474 | 95.7 | 5.85 |
| Cer 38:2;2 | 592.5651 | 73.6 | 2.14 |
| Cer 38:1;2 | 594.5802 | 25.2 | 2.93 |
| Cer 42:2;2 | 648.6256 | 676.3 | 5.15 |
| CerP 34:2;2 | 616.4675 | 5.7 | 4.10 |
| CerP 36:1;2 | 646.5145 | 22.0 | 3.86 |
| CerP 38:2;2 | 672.5293 | 208.5 | 5.04 |
| CerP 40:2;2 | 700.5639 | 14.0 | 0.05 |
| CerP 42:2;2 | 728.5969 | 83.7 | 2.23 |
| CerP 42:1;2 | 730.6142 | 15.6 | 4.56 |
| CerP 44:2;2 | 756.6234 | 33.4 | 4.22 |
| CerP 44:1;2 | 758.6425 | 3.4 | 0.33 |
| SM 36:2;2 | 715.5744 | 5.6 | 0.60 |
| SM 36:1;2 | 717.5880 | 48.6 | 3.47 |

**Supplementary Table 3: Putatively assigned lipids identified within the 1 µm MSI of a cerebellar white matter, having undergone novel pre-staining preparation (cf. Main text Fig. 3a purple-cross).**

IDs are based on accurate mass matching at the $MS^1$ level and thus some misassignments may be present. The ions nonetheless represent sample-related mass features and further support the rich biochemical information obtained.

PC assignments are based off the headgroup loss fragments (-183.066 *m/z*), and SM assignments are based off the loss of a methyl group (-14.016 *m/z*). Thus, all lipid identifications are assigned as $[M+H]^+$ ions, with the exception of PC, SM and cholesterol, which are assigned as $[M-183.066+H]^+$, $[M-14.016+H]^+$and $[M-H_2O+H]^+$, respectively. See the *Methods* for further details regarding source fragmentation of choline containing lipids.

NB: while 203 lipids were identified from the purple-cross region indicated in Main Text Fig. 3a and 3d, a number of the Hex3Cer and Hex2Cer lipids were deemed as 'noisy images' and were manually removed from the total number of putative identifications reported in text (*i.e.,* 191).

| Cerebellum white matter, CV pre-stain, DMACA; 1 µm (203 IDs) | | | |
|---|---|---|---|
| Lipid ID | exp. m/z | Δppm | Averaged abs. intensity (across whole area) |
| PC 28:0 | 495.4397 | 2.128051 | 24.56 |
| PC 32:3 | 545.4579 | 2.745169 | 17.36 |
| PC 32:2 | 547.4689 | 5.790418 | 66.8 |
| PC 34:5 | 569.4584 | 3.389039 | 10.56 |
| PC 34:4 | 571.4721 | 0.034131 | 22.96 |
| PC 34:3 | 573.4844 | 5.880105 | 105.36 |
| PC 34:2 | 575.5008 | 4.441016 | 1020.48 |
| PC 34:1 | 577.5157 | 5.734943 | 6721.28 |
| PC 36:5 | 597.4859 | 3.098468 | 21.68 |
| PC 36:4 | 599.4998 | 5.963871 | 282.56 |
| PC 38:6 | 623.5012 | 3.548752 | 118.4 |
| PC 38:4 | 627.5326 | 3.300894 | 588.48 |
| PC 40:7 | 649.5181 | 1.509502 | 47.44 |
| PC 40:6 | 651.5314 | 5.001389 | 262.88 |
| PC 40:5 | 653.5478 | 3.922293 | 64.16 |
| PC 40:2 | 659.5947 | 3.845735 | 200.32 |
| PC 42:6 | 679.5669 | 1.282972 | 16.48 |
| PC 42:5 | 681.5779 | 5.412037 | 23.12 |
| PC 42:4 | 683.5984 | 1.638662 | 57.04 |
| PC 42:3 | 685.6111 | 2.616381 | 34.48 |
| PC 42:2 | 687.6264 | 3.228232 | 134.64 |
| PC 42:1 | 689.6406 | 5.24748 | 200.48 |
| PC 44:6 | 707.5976 | 0.482742 | 20.72 |
| PC 44:2 | 715.6567 | 4.428374 | 15.6 |
| PE 30:2 | 660.4627 | 4.245141 | 17.76 |
| PE 34:4 | 712.4934 | 3.070153 | 38.72 |
| PE 34:3 | 714.509 | 3.045991 | 76.64 |
| PE 34:2 | 716.5196 | 4.019884 | 158.88 |
| PE 36:5 | 738.5033 | 4.757595 | 31.12 |
| PE 36:4 | 740.5185 | 5.399095 | 107.28 |
| PE 36:3 | 742.5354 | 3.631678 | 500.64 |

| | | | |
|---|---|---|---|
| PE 36:2 | 744.5504 | 4.495488 | 1028.8 |
| PE 38:6 | 764.518 | 5.847073 | 124.8 |
| PE 38:5 | 766.5352 | 3.860747 | 225.04 |
| PE 38:4 | 768.55 | 4.95985 | 516.32 |
| PE 40:6 | 792.55 | 4.792473 | 311.2 |
| PE 40:3 | 798.5963 | 5.538772 | 54.8 |
| PE 40:2 | 800.6153 | 1.336801 | 37.92 |
| PE 42:3 | 826.6369 | 5.909448 | 82.96 |
| PE 42:2 | 828.6481 | 0.488308 | 125.44 |
| PE 42:1 | 830.6642 | 0.984052 | 38.48 |
| PE 42:0 | 832.681 | 2.367984 | 11.6 |
| PE 44:2 | 856.6809 | 2.28675 | 23.36 |
| PS 32:3 | 730.4645 | 1.214495 | 37.04 |
| PS 32:2 | 732.4833 | 3.179554 | 19.68 |
| PS 32:1 | 734.5004 | 5.146107 | 27.68 |
| PS 34:5 | 754.4625 | 3.779393 | 15.52 |
| PS 34:3 | 758.4983 | 2.189647 | 16.72 |
| PS 34:2 | 760.511 | 1.714064 | 23.6 |
| PS 36:5 | 782.5013 | 5.921875 | 19.12 |
| PS 36:3 | 786.5258 | 2.697374 | 16.88 |
| PS 36:2 | 788.5419 | 2.138773 | 22.32 |
| PS 36:1 | 790.5553 | 5.046139 | 83.04 |
| PS 38:4 | 812.5403 | 4.021722 | 15.28 |
| PS 38:2 | 816.5724 | 3.090019 | 15.76 |
| PS 40:6 | 836.5452 | 1.903407 | 15.12 |
| PS 40:5 | 838.5637 | 5.303086 | 15.04 |
| PS 40:4 | 840.5703 | 5.519422 | 20.72 |
| PS 40:3 | 842.5943 | 4.408203 | 9.84 |
| PS 40:2 | 844.6063 | 0.058494 | 30.32 |
| PS 40:0 | 848.6404 | 3.418164 | 94.56 |
| PS 42:2 | 872.6421 | 5.297775 | 72.24 |
| PS 42:1 | 874.6559 | 3.124131 | 90.32 |
| PS 44:11 | 882.5324 | 5.069693 | 5.52 |
| PS 44:10 | 884.5425 | 1.277036 | 9.84 |
| PS 44:7 | 890.5912 | 0.744809 | 10.8 |
| PS 44:6 | 892.6018 | 4.971978 | 21.52 |
| PI 28:0 | 755.4742 | 4.844271 | 19.76 |
| PI 34:6 | 827.4681 | 2.90476 | 6.8 |
| PI 36:8 | 851.4668 | 4.310603 | 7.36 |
| PI 36:7 | 853.4829 | 3.771326 | 8.08 |
| PI 38:8 | 879.4973 | 5.075889 | 4.8 |
| PI 38:7 | 881.5225 | 5.736806 | 3.12 |
| PI 38:6 | 883.5303 | 3.222099 | 9.12 |
| PI 38:1 | 893.6155 | 4.581241 | 14.24 |
| PI 40:5 | 913.5751 | 5.477942 | 4.16 |
| PI 42:4 | 943.6219 | 5.446181 | 10 |

| | | | |
|---|---|---|---|
| PI 42:2 | 947.655 | 3.480711 | 4.56 |
| PI 42:0 | 951.6911 | 1.581103 | 4.88 |
| PI 44:11 | 957.5525 | 3.876957 | 2.32 |
| PI 44:9 | 961.5826 | 2.692452 | 2.88 |
| PI 44:8 | 963.5945 | 1.299555 | 5.52 |
| PI 44:6 | 967.6307 | 3.825112 | 2.08 |
| PI 44:3 | 973.6779 | 4.011758 | 2.8 |
| PI 44:1 | 977.7026 | 2.739665 | 8.24 |
| PG 30:0 | 695.4823 | 4.927781 | 10.16 |
| PG 38:7 | 793.4976 | 4.746338 | 11.36 |
| PG 38:0 | 807.6151 | 5.108011 | 13.52 |
| PG 40:8 | 819.5214 | 5.345633 | 18.08 |
| PG 42:10 | 843.5216 | 5.425731 | 5.36 |
| PG 42:2 | 859.6373 | 5.77871 | 9.52 |
| PG 44:9 | 873.5633 | 0.864247 | 6.72 |
| PG 44:4 | 883.6407 | 1.758266 | 11.52 |
| PA 34:5 | 667.4307 | 3.884299 | 11.44 |
| PA 36:5 | 695.4615 | 4.54567 | 19.28 |
| PA 36:4 | 697.4778 | 3.632838 | 45.12 |
| PA 36:1 | 703.523 | 5.976711 | 264.72 |
| PA 38:8 | 717.4481 | 1.23724 | 27.84 |
| PA 38:7 | 719.467 | 3.233059 | 30.48 |
| PA 38:6 | 721.4771 | 4.447659 | 51.84 |
| PA 38:5 | 723.4928 | 4.323201 | 84.24 |
| PA 40:8 | 745.478 | 3.093092 | 28.32 |
| PA 40:4 | 753.5426 | 0.338627 | 93.68 |
| PA 42:9 | 771.4992 | 4.182391 | 12.72 |
| PA 42:8 | 773.5115 | 0.062836 | 36.4 |
| PA 42:7 | 775.5253 | 2.517607 | 75.52 |
| PA 42:6 | 777.5443 | 1.782807 | 67.28 |
| PA 42:5 | 779.5607 | 2.803441 | 30.4 |
| PA 44:11 | 795.4998 | 4.838346 | 10.32 |
| PA 44:10 | 797.5149 | 4.214162 | 18.88 |
| PA 44:9 | 799.5272 | 0.022131 | 14.4 |
| PA 44:8 | 801.5446 | 2.097312 | 29.52 |
| PA 44:7 | 803.563 | 5.540566 | 21.28 |
| PA 44:6 | 805.5744 | 0.275911 | 34.08 |
| PA 44:5 | 807.5868 | 3.728055 | 17.44 |
| PA 44:4 | 809.6043 | 1.502581 | 15.52 |
| PC-O 28:1 | 479.4457 | 0.276853 | 16.8 |
| PC-O 28:0 | 481.4636 | 4.410329 | 7.2 |
| PC-O 30:0 | 509.4902 | 5.055501 | 18.24 |
| PC-O 34:5 | 555.4762 | 1.830741 | 10.96 |
| PC-O 36:2 | 589.5536 | 3.065569 | 90.48 |
| PC-O 36:0 | 593.5851 | 2.659778 | 77.84 |
| PC-O 38:3 | 615.5712 | 0.243087 | 22.32 |

| | | | |
|---|---|---|---|
| PC-O 38:2 | 617.5874 | 1.152295 | 29.2 |
| PC-O 42:6 | 665.5878 | 1.688755 | 44.4 |
| PC-O 44:7 | 691.5985 | 5.565779 | 24.32 |
| PC-O 44:6 | 693.6209 | 4.208473 | 18.56 |
| PC-O 44:5 | 695.6319 | 2.572429 | 10.48 |
| PC-O 44:0 | 705.7144 | 3.500003 | 1.76 |
| PE-O 28:2 | 618.4496 | 0.404326 | 18.8 |
| PE-O 34:5 | 696.4985 | 3.17922 | 48.72 |
| PE-O 34:3 | 700.5289 | 1.892745 | 119.76 |
| PE-O 36:5 | 724.5281 | 0.790412 | 44.24 |
| PE-O 36:4 | 726.5415 | 2.367497 | 65.12 |
| PE-O 36:3 | 728.5568 | 2.828628 | 348.24 |
| PE-O 36:2 | 730.5704 | 5.635495 | 285.04 |
| PE-O 36:1 | 732.5896 | 0.830319 | 47.68 |
| PE-O 38:6 | 750.542 | 1.64295 | 35.2 |
| PE-O 38:5 | 752.5561 | 3.632177 | 31.84 |
| PE-O 38:4 | 754.5719 | 3.447582 | 43.52 |
| PE-O 40:5 | 780.5865 | 4.678765 | 56.48 |
| PE-O 40:4 | 782.6069 | 1.435745 | 24.56 |
| PE-O 40:3 | 784.6208 | 0.816188 | 27.44 |
| PE-O 40:2 | 786.6399 | 3.531389 | 12.64 |
| PE-O 42:5 | 808.6252 | 4.663567 | 21.04 |
| PE-O 42:4 | 810.6372 | 0.065348 | 48.24 |
| PE-O 42:2 | 814.672 | 4.38643 | 383.28 |
| PE-O 42:1 | 816.6826 | 1.755331 | 95.04 |
| PE-O 44:4 | 838.6727 | 5.141827 | 16.4 |
| PE-O 44:3 | 840.688 | 4.631411 | 24.32 |
| PE-O 44:2 | 842.7038 | 4.869651 | 37.28 |
| HexCer 34:1;2 | 700.5709 | 1.810089 | 20.64 |
| HexCer 34:0;2 | 702.588 | 0.17197 | 18.08 |
| HexCer 36:2;2 | 726.5851 | 3.789362 | 436.48 |
| HexCer 40:2;2 | 782.6461 | 5.586228 | 3617.84 |
| HexCer 40:0;2 | 786.6792 | 3.198386 | 56.4 |
| HexCer 42:2;2 | 810.6777 | 4.984899 | 4796.96 |
| HexCer 44:2;2 | 838.7105 | 3.068513 | 33.2 |
| Hex2Cer 28:0;2 | 780.5475 | 0.919181 | 38.32 |
| Hex2Cer 32:1;2 | 834.5942 | 0.517076 | 16.48 |
| Hex2Cer 34:2;2 | 860.6135 | 4.843466 | 15.44 |
| Hex2Cer 34:1;2 | 862.6254 | 0.461735 | 31.52 |
| Hex2Cer 34:0;2 | 864.6377 | 3.460044 | 26 |
| Hex2Cer 36:1;2 | 890.6536 | 3.087758 | 60.08 |

| | | | |
|---|---|---|---|
| Hex2Cer 38:2;2 | 916.6682 | 4.154555 | 6.72 |
| Hex2Cer 38:1;2 | 918.6825 | 5.619391 | 8.96 |
| Hex2Cer 38:0;2 | 920.7012 | 2.270364 | 3.28 |
| Hex2Cer 40:2;2 | 944.7029 | 0.354239 | 3.44 |
| Hex2Cer 40:1;2 | 946.7173 | 1.71483 | 3.44 |
| Hex2Cer 42:0;2 | 976.7693 | 3.532504 | 2.4 |
| Hex2Cer 44:2;2 | 1000.771 | 4.65103 | 4.88 |
| Hex3Cer 28:2;2 | 938.5635 | 5.097867 | 4.96 |
| Hex3Cer 28:0;2 | 942.6033 | 3.939786 | 5.52 |
| Hex3Cer 30:0;2 | 970.635 | 4.248645 | 4.8 |
| Hex3Cer 32:1;2 | 996.6509 | 4.319554 | 2.4 |
| Hex3Cer 34:0;2 | 1026.695 | 1.780849 | 3.6 |
| Hex3Cer 36:0;2 | 1054.72 | 4.662726 | 1.04 |
| Hex3Cer 42:2;2 | 1134.787 | 0.125533 | 2.48 |
| Hex3Cer 42:0;2 | 1138.817 | 1.303065 | 1.84 |
| Hex3Cer 44:2;2 | 1162.822 | 3.216307 | 1.28 |
| Hex3Cer 44:1;2 | 1164.83 | 3.700832 | 0.96 |
| Cer 28:2;2 | 452.4086 | 2.708517 | 58.4 |
| Cer 30:2;2 | 480.4392 | 3.957727 | 14.16 |
| Cer 34:2;2 | 536.5049 | 2.195596 | 13.84 |
| Cer 36:2;2 | 564.5369 | 3.257532 | 166.72 |
| Cer 36:1;2 | 566.5502 | 0.845083 | 54.96 |
| Cer 38:2;2 | 592.568 | 2.862815 | 103.76 |
| Cer 38:1;2 | 594.5832 | 2.073845 | 50.64 |
| Cer 40:1;2 | 622.6102 | 4.924209 | 134.96 |
| Cer 42:2;2 | 648.6256 | 5.145926 | 1867.12 |
| CerP 30:1;2 | 562.421 | 3.741444 | 27.68 |
| CerP 34:2;2 | 616.4737 | 5.904761 | 10.48 |
| CerP 36:2;2 | 644.5037 | 3.7155 | 20.88 |
| CerP 38:2;2 | 672.5293 | 5.040362 | 21.12 |
| CerP 40:0;2 | 704.5932 | 2.917945 | 18.16 |
| CerP 42:2;2 | 728.5932 | 2.768453 | 96.4 |
| CerP 42:1;2 | 730.6106 | 0.437557 | 42.24 |

| | | | |
|---|---|---|---|
| CerP 42:0;2 | 732.6262 | 0.495342 | 10.8 |
| CerP 44:2;2 | 756.6271 | 0.782937 | 35.36 |
| CerP 44:1;2 | 758.6425 | 0.328443 | 9.44 |
| SM 32:1;2 | 661.5302 | 3.466898 | 15.68 |
| SM 34:0;2 | 691.5778 | 4.22522 | 15.2 |
| SM 40:0;2 | 775.6649 | 4.980734 | 15.12 |

**Supplementary Table 4: Putatively assigned lipids identified within the 1 µm MSI of a cerebellar grey matter, having undergone novel pre-staining preparation (cf. Main text Fig. 3a blue-cross).**

IDs are based on accurate mass matching at the MS[1] level and thus some misassignments may be present. The ions nonetheless represent sample-related mass features and further support the rich biochemical information obtained.

PC assignments are based off the headgroup loss fragments (-183.066 *m/z*), and SM assignments are based off the loss of a methyl group (-14.016 *m/z*). Thus, all lipid identifications are assigned as $[M+H]^+$ ions, with the exception of PC, SM and cholesterol, which are assigned as $[M-183.066+H]^+$, $[M-14.016+H]^+$and $[M-H_2O+H]^+$, respectively. See the *Methods* for further details regarding source fragmentation of choline containing lipids.

NB: while 139 lipids were identified from the blue-cross region indicated in Main Text Fig. 3a and 3d, a number of the Hex3Cer and Hex2Cer lipids were deemed as 'noisy images' and were manually removed from the total number of putative identifications reported in text (*i.e.,* 133).

| Cerebellum grey matter, CV pre-stain, DMACA; 1 µm (139 IDs) | | | |
|---|---|---|---|
| Lipid ID | exp. m/z | Δppm | Averaged abs. intensity (across whole area) |
| PC 28:0 | 495.4397 | 2.128051 | 11.6 |
| PC 30:1 | 521.4538 | 5.03509 | 46 |
| PC 30:0 | 523.4705 | 3.008261 | 64.48 |
| PC 32:2 | 547.4689 | 5.790418 | 61.6 |
| PC 32:1 | 549.4846 | 5.770713 | 418.56 |
| PC 34:4 | 571.4721 | 0.034131 | 13.44 |
| PC 34:3 | 573.4844 | 5.880105 | 136.8 |
| PC 34:2 | 575.5008 | 4.441016 | 1329.76 |
| PC 34:1 | 577.5157 | 5.734943 | 5596.72 |
| PC 36:4 | 599.4998 | 5.963871 | 85.92 |
| PC 36:2 | 603.533 | 2.856995 | 848.32 |
| PC 38:7 | 621.4843 | 5.529896 | 21.36 |
| PC 38:3 | 629.5471 | 5.168025 | 18.56 |
| PC 40:6 | 651.5314 | 5.001389 | 112.48 |
| PC 40:3 | 657.5795 | 3.286331 | 3.44 |
| PC 40:1 | 661.6096 | 5.066689 | 9.76 |
| PC 42:5 | 681.5779 | 5.412037 | 5.2 |
| PC 42:2 | 687.6264 | 3.228232 | 3.6 |
| PC 42:1 | 689.6441 | 0.247495 | 7.2 |
| PC 44:4 | 711.6318 | 4.481417 | 3.92 |
| PE 32:1 | 690.5032 | 5.251754 | 16.4 |
| PE 32:0 | 692.5224 | 0.088065 | 35.2 |
| PE 34:6 | 708.4579 | 2.825922 | 13.92 |
| PE 34:2 | 716.5196 | 4.019884 | 23.2 |
| PE 34:0 | 720.5506 | 4.455268 | 126.96 |
| PE 36:4 | 740.5185 | 5.399095 | 24.16 |
| PE 36:3 | 742.5354 | 3.631678 | 34.08 |
| PE 36:2 | 744.5542 | 0.504501 | 51.76 |
| PE 36:1 | 746.5672 | 3.03039 | 168.72 |
| PE 38:7 | 762.5062 | 0.882362 | 10.56 |
| PE 38:5 | 766.5352 | 3.860747 | 8.56 |

| | | | |
|---|---|---|---|
| PE 38:4 | 768.5538 | 0.040137 | 19.04 |
| PE 38:3 | 770.5662 | 4.180626 | 15.84 |
| PE 38:2 | 772.5839 | 1.559008 | 15.6 |
| PE 40:8 | 788.5183 | 5.342158 | 10.56 |
| PE 40:6 | 792.55 | 4.792473 | 25.52 |
| PE 40:1 | 802.6274 | 5.79635 | 2.88 |
| PE 42:6 | 820.5834 | 2.036023 | 6.4 |
| PE 42:0 | 832.681 | 2.367984 | 3.6 |
| PE 44:12 | 836.5243 | 2.162429 | 5.92 |
| PE 44:5 | 850.6286 | 4.086394 | 2.64 |
| PE 44:4 | 852.6427 | 5.87502 | 1.12 |
| PE 44:3 | 854.6658 | 2.924791 | 3.52 |
| PE 44:2 | 856.6767 | 2.713248 | 2.32 |
| PS 32:3 | 730.4681 | 3.78551 | 14.96 |
| PS 32:1 | 734.4968 | 0.146095 | 7.92 |
| PS 32:0 | 736.5121 | 0.356364 | 16.24 |
| PS 34:3 | 758.4945 | 2.810349 | 7.04 |
| PS 36:5 | 782.4974 | 0.92186 | 6.96 |
| PS 38:5 | 810.5237 | 5.272184 | 5.52 |
| PS 38:3 | 814.5579 | 1.617532 | 5.52 |
| PS 40:5 | 838.5637 | 5.303086 | 4.72 |
| PS 40:4 | 840.5745 | 0.519438 | 5.2 |
| PS 44:5 | 894.6169 | 5.589463 | 6.64 |
| PI 30:2 | 779.475 | 5.726334 | 8.72 |
| PI 34:1 | 837.5497 | 1.07789 | 3.76 |
| PI 36:7 | 853.4829 | 3.771326 | 4.64 |
| PI 36:0 | 867.5911 | 5.283938 | 5.76 |
| PI 38:0 | 895.6239 | 3.501956 | 4.32 |
| PI 40:6 | 911.5628 | 1.719744 | 4.96 |
| PI 40:5 | 913.5751 | 5.477942 | 1.92 |
| PI 42:5 | 941.6094 | 2.094463 | 4.32 |
| PI 42:2 | 947.655 | 3.480711 | 1.36 |
| PI 44:5 | 969.6455 | 2.905393 | 2.32 |
| PI 44:4 | 971.6596 | 1.306908 | 1.52 |
| PG 28:2 | 663.4249 | 2.585611 | 11.76 |
| PG 36:7 | 765.4743 | 5.413069 | 4.72 |
| PG 40:5 | 825.563 | 1.242219 | 3.28 |
| PG 40:4 | 827.5757 | 4.810158 | 3.84 |
| PG 44:11 | 869.5324 | 0.408509 | 7.12 |
| PG 44:9 | 873.5676 | 4.13576 | 3.84 |
| PG 44:7 | 877.5997 | 4.976392 | 1.76 |
| PG 44:3 | 885.6533 | 5.231396 | 1.12 |
| PG 44:1 | 889.6877 | 1.662275 | 0.64 |
| PG 44:0 | 891.7096 | 5.33318 | 1.36 |
| PA 30:2 | 617.4145 | 5.104455 | 8.96 |
| PA 34:6 | 665.4181 | 0.68799 | 12.24 |

| | | | |
|---|---|---|---|
| PA 34:5 | 667.4341 | 1.115692 | 9.6 |
| PA 36:6 | 693.451 | 2.94154 | 11.6 |
| PA 36:5 | 695.4615 | 4.54567 | 17.92 |
| PA 36:4 | 697.4778 | 3.632838 | 39.04 |
| PA 36:2 | 701.5103 | 1.800513 | 125.04 |
| PA 38:4 | 725.5105 | 1.436998 | 23.76 |
| PA 38:3 | 727.5266 | 0.832101 | 18 |
| PA 40:8 | 745.478 | 3.093092 | 27.76 |
| PA 40:7 | 747.4935 | 3.272501 | 51.44 |
| PA 42:8 | 773.5077 | 5.062822 | 14.88 |
| PA 42:7 | 775.5253 | 2.517607 | 25.2 |
| PA 42:5 | 779.5607 | 2.803441 | 12.56 |
| PA 42:4 | 781.5746 | 0.509577 | 4.96 |
| PA 44:10 | 797.511 | 0.785845 | 7.92 |
| PA 44:9 | 799.5232 | 5.022117 | 4.88 |
| PA 44:5 | 807.5868 | 3.728055 | 6.96 |
| PA 44:4 | 809.6083 | 3.497423 | 6.64 |
| PA 44:0 | 817.6673 | 0.922267 | 0.64 |
| PC-O 32:2 | 533.4902 | 4.92108 | 5.04 |
| PC-O 32:0 | 537.5225 | 2.94436 | 10.08 |
| PC-O 34:6 | 553.4607 | 1.54805 | 9.84 |
| PC-O 34:0 | 565.5539 | 2.618427 | 12.08 |
| PC-O 38:3 | 615.5712 | 0.243087 | 3.28 |
| PC-O 38:2 | 617.5874 | 1.152295 | 2 |
| PC-O 40:4 | 641.5843 | 3.721119 | 2.16 |
| PC-O 42:4 | 669.6168 | 1.80604 | 4.24 |
| PC-O 44:7 | 691.6054 | 4.434212 | 6.56 |
| PE-O 28:2 | 618.4465 | 4.595662 | 11.04 |
| PE-O 32:0 | 678.5449 | 2.4483 | 2.56 |
| PE-O 34:5 | 696.495 | 1.820782 | 6.16 |
| PE-O 36:2 | 730.5777 | 4.364495 | 3.12 |
| PE-O 42:7 | 804.5922 | 2.531782 | 8.16 |
| PE-O 42:2 | 814.6638 | 5.613561 | 1.92 |
| PE-O 44:10 | 826.5749 | 0.489385 | 2.64 |
| PE-O 44:3 | 840.6838 | 0.368598 | 1.84 |
| HexCer 38:1;2 | 756.6309 | 5.110081 | 12.32 |
| HexCer 40:2;2 | 782.65 | 0.586245 | 4.8 |
| HexCer 40:1;2 | 784.664 | 2.691692 | 2.4 |
| HexCer 42:1;2 | 812.6988 | 1.736127 | 3.68 |
| Hex2Cer 30:2;2 | 804.548 | 1.47127 | 7.28 |
| Hex2Cer 30:1;2 | 806.5659 | 4.28004 | 7.76 |
| Hex2Cer 32:1;2 | 834.5942 | 0.517076 | 4.32 |
| Hex2Cer 34:2;2 | 860.6049 | 5.15653 | 2.88 |

| | | | |
|---|---|---|---|
| Hex2Cer 38:2;2 | 916.6727 | 0.845435 | 2.64 |
| Hex2Cer 40:2;2 | 944.7077 | 4.64577 | 3.92 |
| Hex2Cer 40:0;2 | 948.7312 | 3.542381 | 1.36 |
| Hex3Cer 30:2;2 | 966.5959 | 3.823247 | 1.28 |
| Hex3Cer 32:0;2 | 998.6611 | 1.070824 | 4.24 |
| Hex3Cer 44:2;2 | 1162.822 | 3.216307 | 0.64 |
| Cer 36:1;2 | 566.553 | 4.154924 | 12.88 |
| Cer 36:0;2 | 568.565 | 2.289968 | 5.92 |
| Cer 38:0;2 | 596.5963 | 2.222757 | 2 |
| Cer 42:2;2 | 648.6256 | 5.145926 | 3.44 |
| Cer 44:2;2 | 676.6611 | 1.366528 | 1.92 |
| CerP 28:2;2 | 532.3737 | 4.600238 | 14.48 |
| CerP 40:2;2 | 700.5604 | 5.045335 | 10.96 |
| CerP 42:1;2 | 730.6106 | 0.437557 | 1.68 |
| CerP 42:0;2 | 732.6225 | 5.495325 | 1.28 |
| CerP 44:2;2 | 756.6309 | 5.782952 | 12.32 |
| SM 32:1;2 | 661.5302 | 3.466898 | 6.96 |
| SM 36:1;2 | 717.588 | 3.469497 | 13.28 |
| SM 44:2;2 | 827.704 | 4.730392 | 2.16 |
